# Neuromodulators control neuronal dynamics through feature space reshaping

**DOI:** 10.1101/2025.05.19.654470

**Authors:** Domenico Guarino, Ilaria Carannante, Alain Destexhe

## Abstract

The activity of neurons in the central nervous system changes drastically across different states like asleep or awake, attentive or drowsy, executing motor commands or resting. Neuromodulation supports these changes by affecting neuronal excitability. Much work has been devoted to understanding the effects of neuromodulators on various neurons. However, we still lack a cohesive picture of how neuromodulation affects neuronal dynamics. Here, we analysed electrophysiological data from published papers and extracted features for seven types of neurons from five areas of the human and rodent central nervous system, under the effect of five neuromodulators. We studied neuromodulation dynamics using the widespread, simple, yet biologically accurate, Adaptive Exponential Integrate-and-Fire model. We have found that different neuromodulators remap the feature space of neurons into non-overlapping clusters. This work organises our knowledge about neuromodulation in a compact format, useful for modellers and theoreticians.

## Introduction

Neuromodulation continuously influences our nervous system, dynamically shaping neuronal activity to regulate various cognitive, emotional, and behavioural functions (***Marder, 2012***). Neuromodulators exert a variety of effects on brain cells, including changing neuronal excitability, shaping short and long-term synaptic plasticity (***Nadim and Bucher, 2014***) and altering astrocytic calcium signalling (***Stevenson et al., 2020***). In this study, we focus on the neuromodulators’ ability to alter the excitability and spiking activity of neurons through mechanisms endogenous to the nervous system (***Kaczmarek et al., 1987; Nadim and Bucher, 2014***).

The internal and external concentration of ions determines the excitability of neurons. For a typical neuron at rest, the concentration of sodium (*Na*^+^), chloride (*Cl*^−^), and calcium (*Ca*^2+^) is greater outside the neuron, whereas the concentration of potassium (*K*^+^) and other anions is greater inside. When a neurotransmitter or other substance binds to a receptor on the membrane, it may change the permeability to some ionic species by opening or closing ionic channels (***Hille, 2001***, p.38, 72). As a consequence, the internal and external concentrations will change. For example, an external reduction or internal augmentation in the *Na*^+^ and *Ca*^2+^ will lead to a more positively charged, i.e. excited, membrane.

There are hundreds of specific receptors for molecules capable of altering membrane permeabilities. These are usually divided into: amino acids (such as Glutamate and GABA), peptides (like Oxytocin and other endorphins), monoamines (including Dopamine, Adrenalin, Serotonin, Histamine and Norepinephrine), purines (Adenosine triphosphate, Adenosine), gasotransmitters (Carbon monoxide, Nitric oxide), and Acetylcholine (***Epelbaum, 1995***). Another common manner of dividing these molecules is into neurotransmitters (such as Glutamate and GABA) and neuromodulators (such as Acetylcholine and Dopamine). Neuromodulators differ from neurotransmitters in that (i) they are typically expressed by specific groups of neurons, (ii) project diffusely throughout the nervous system, (iii) usually have longer-lasting effects, and (iv) they modulate postsynaptic neurons by altering their responses to neurotransmitters (***Epelbaum, 1995; Hille, 2001***, p.212).

There have been several modeling works detailing the interaction between specific neuromodulators, their target channels, and their effects on multicompartmental membrane models and microcircuits (***Lindroos et al., 2018; Frost Nylen et al., 2021***). In addition, several frameworks exist for optimising the parameter space of detailed Hodgkin-Huxley models (***Achard and De Schutter, 2006; Svensson et al., 2012; Van Geit et al., 2016***). However, the main problem is that we are still lacking a cohesive general picture of how different families of neuromodulators affect neuronal dynamics. Considering detailed ion channel-based models would lead to the impossibility of comparing different neurons expressing different types and distributions of channels. To consolidate our understanding of neuromodulation dynamics, and foster the desired cohesion at a manageable scale, we adopted, here, the simple yet dynamically rich and conductance-based Adaptive Exponential Integrate-and-Fire (AdEx) model (***Brette and Gerstner, 2005; Touboul and Brette, 2008***).

First, we aggregated different sources in the literature concerning seven types of neurons from five areas of the human and rodent central nervous system under the effect of five types of neuromodulators. Second, we extracted key features from electrophysiological recordings illustrated in published papers, and we conducted a parameter space exploration to fit the AdEx model parameters. Third, we analysed how neuromodulation influences the parameters of the control models and applied a dimensionality reduction technique, specifically principal component analysis (PCA), to visualise the parameter space and evaluate the formation of independent clusters. Finally, we studied the phase plane of this two-dimensional system.

This work aims to help bridge the gap between the electrophysiological and computational communities by considering several neuromodulators and neuron types, in a simple computational neuron model, and providing a framework for representing and interpreting the effects of neuromodulation on neuron excitability in a concise, reproducible and accessible manner.

## Results

The literature provides a range of effects that neuromodulators have on different neuron types. However, there are two main issues when facing such a range of effects. Its variety limits the ability to compare datasets, and the actual available data is often limited to a few electrophysiological traces. To build a comparable dataset, we identified a minimal number of features to describe electrophysiological traces reported in published papers from neurons in control and under the effects of neuromodulators. These include time to first, second, third, and last spike (red bars 1, 2, 3, 4 in Fig.1A and B), inverse of the first and last interspike interval (red bars 5 and 6, in Fig.1A and B), and firing frequency. When modelling electrophysiological traces showing strong adaptation, we have included the value of the voltage and the end of the stimulus. We chose these features because they are readily accessible, capable of capturing a wide range of electrophysiological behaviours, and compatible with the level of detail we can retain when extracting traces from different publications.

We extracted and collected the key features of seven types of neurons – Pyramidal, direct and indirect Striatal projection, Granule, Relay, Reticular, Purkinje – from five areas of the human and rodent central nervous system – Cortex, Striatum, Hippocampus, Thalamus and Cerebellum – under the effect of five types of neuromodulators – Acetylcholine, Norepinephrine, Serotonin, Histamine, and Dopamine (Table 1).

**Table 1.** Dataset of electrophysiological recordings across brain regions, species, and neuromodulators. * Muscarine, a selective agonist of Acetylcholine was bath applied. ** Oxo-M, a muscarinic agonist was bath applied. *** Isoproterenol, a *β*-adrenergic agonist was bath applied.

| Input Region | Species | Condition | Publication | Figure |
| --- | --- | --- | --- | --- |
| Cortex | Human | Control | *McCormick and Williamson, 1989* | Fig. 1A |
| Cortex | Human | Control | *McCormick and Williamson, 1989* | Fig. 1B |
| Cortex | Human | Control | *McCormick and Williamson, 1989* | Fig. 1D |
| Cortex | Human | Histamine | *McCormick and Williamson, 1989* | Fig. 1A |
| Cortex | Human | Methacoline | *McCormick and Williamson, 1989* | Fig. 1B |
| Cortex | Human | Norepinephrine | *McCormick and Williamson, 1989* | Fig. 1C |
| Cortex | Human | Serotonin | *McCormick and Williamson, 1989* | Fig. 1D |
| Cortex | Rat | Control | *Onn et al., 2006* | Fig. 3a1 |
| Cortex | Rat | Dopamine | *Onn et al., 2006* | Fig. 3a2 |
| Striatum | Mouse | D1_control | *Planert et al., 2013* | Fig. 4C |
| Striatum | Mouse | D2_control | *Planert et al., 2013* | Fig. 4C |
| Striatum | Mouse | D1_dopamine | *Planert et al., 2013* | Fig. 4C |
| Striatum | Mouse | D2_dopamine | *Planert et al., 2013* | Fig. 4C |
| Striatum | Rat | Control | *Perez-Rosello et al., 2005* | Fig. 8A |
| Striatum | Rat | Acetylcholine* | *Perez-Rosello et al., 2005* | Fig. 8B |
| Hippocampus | Mouse | Control | *Carver and Shapiro, 2019* | Fig. 5A (60pA) |
| Hippocampus | Mouse | Control | *Carver and Shapiro, 2019* | Fig. 5A (100pA) |
| Hippocampus | Mouse | Control | *Carver and Shapiro, 2019* | Fig. 5A (140pA) |
| Hippocampus | Mouse | Acetylcholine** | *Carver and Shapiro, 2019* | Fig. 5A (140pA) |
| Hippocampus | Rat | Histamine | *Greene and Haas, 1990* | Fig. 3 |
| Hippocampus | Rat | Norepinephrine*** | *Haas and Rose, 1987* | Fig. 2A |
| Thalamus | Rat | TPS Control | *Govindaiah and Cox, 2006* | Fig. 5Aa |
| Thalamus | Rat | TPS Orexin-B | *Govindaiah and Cox, 2006* | Fig. 5Ac |
| Thalamus | Rat | TPS Norepinephrine | *McCormick, 1989* | Fig. 2B |
| Thalamus | Rat | TPS Acetylcholine | *McCormick, 1989* | Fig. 4A |
| Thalamus | Rat | TRN Control | *McCormick and Prince, 1986* | Fig. 3A |
| Thalamus | Rat | TRN Acetylcholine | *McCormick and Prince, 1986* | Fig. 3A |
| Cerebellum | Rat | Control | *Fleming and Hull, 2019* | Fig. 1F |
| Cerebellum | Rat | Norepinephrine | *Fleming and Hull, 2019* | Fig. 1F |
| Cerebellum | Rat | Acetylcholine* | *Watkins and Mathie, 1996* | Fig. 6C |

We use the Adaptive Exponential Integrate-and-Fire model (AdEx), introduced by ***Brette and Gerstner, 2005***, theoretically characterised by ***Touboul and Brette, 2008***, and experimentally characterised for the cortex by ***Naud et al., 2008***. This model is powerful enough to capture the rich and diverse dynamic of neurons under different neuromodulators. Despite the small number of parameters, it retains a clear biological meaning, and its computational efficiency makes it suitable for large-scale network simulations.

The AdEx describes the evolution of the membrane potential *V* (*t*) of a single compartment of a neuron membrane. It consists of a system of two differential equations:

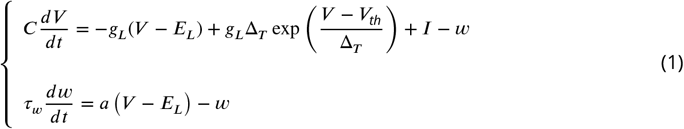

When the potential goes beyond *V*_*th*_, the exponential term generates an upswing of the membrane potential that mimics the exponential activation of *Na*^+^ channels in a Hodgkin–Huxley-type neuron model, leading to the initiation of an action potential. At a spike detection threshold (e.g. 0 mV), a series of reset conditions is triggered:

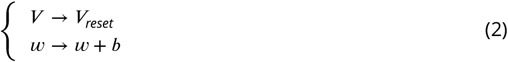

The increase of the adaptation variable *w* by a value *b* leads to accumulation during a spike train. There are nine parameters required to define the evolution of the membrane potential (*V*) and the adaptation current (*w*), divided into *scaling parameters*: *C* membrane capacitance, *g*_*L*_ leak conductance, *E*_*L*_ leak equilibrium potential, Δ_*T*_ exponential slope factor, *V*_*T*_ effective threshold potential, and *bifurcation parameters*: *a* subthreshold adaptation conductance, *b* spike-triggered adaptation, *τ*_*w*_ adaptation time constant, and *V*_*reset*_ after-spike reset potential. Modifying the bifurcation parameters yields qualitative changes in the behaviour of the system.

We created a parameters search procedure, whose inputs are the extracted key features and what we expect to be the interval of variability of each parameter of the AdEx. Briefly, for each parameter, the algorithm samples a value within the given bounds and creates a list of models (each containing nine parameter values). The models are then simulated, and the corresponding features are extracted. Afterwards, an error function quantifies the difference between the data-derived and the model-derived features. Finally, the models are ranked in ascending order of error. We chose the top 16 fits for display and discussion (Fig. 1A and B, grey traces, see Supplementary Fig. 2 and Method for details).

**Figure 1.**
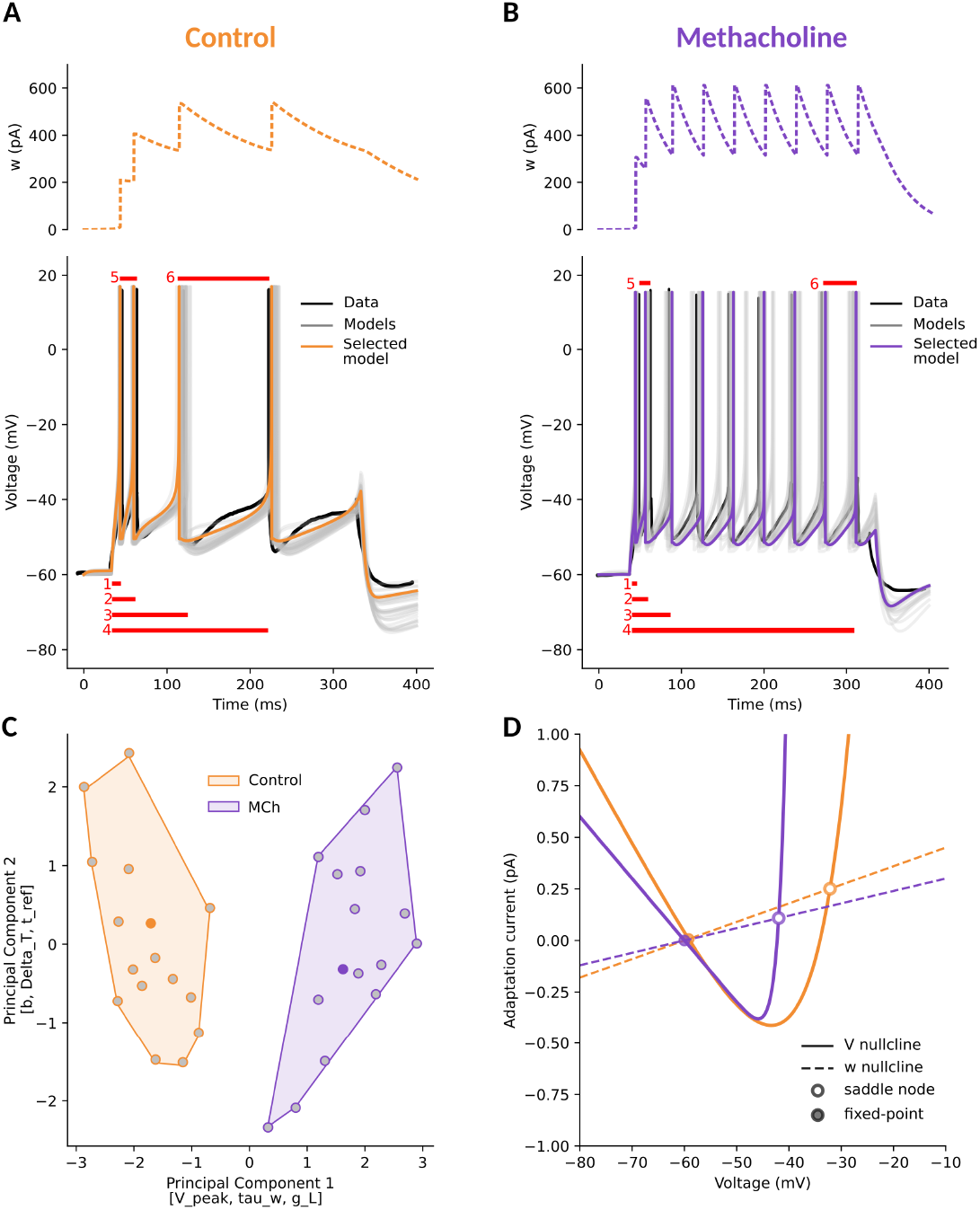
Neuromodulators control the feature space of neurons. **A**. Cortical pyramidal neuron in control condition. Electrophysiology data (black), multiple fitted AdEx models (grey), and one selected model (orange, same colour throughout the paper) are presented together (Voltage traces, solid, on bottom, adaptation current, dashed, on top). The features measured over the traces were: time to first, second, third, and last spike (red bars 1, 2, 3, 4), inverse of the first and last interspike interval (bars 5 and 6), and firing frequency. **B**. The same pyramidal neuron in Methacholine (MCh) bath application. The voltage and adaptation traces are appreciably different. The same features as in A were measured. **C**. To study the relationship between the two conditions, we applied a dimensional reduction technique (PCA) to the parameters of fitted (grey points) and selected (centroid, same colour as in the respective panels) models. In this space, where the top three features contributing to the first principal component were [`V_peak, tau_w, g_L`] and the top contributors of the second principal component were [`b, Delta_T, t_ref`], the control and modulated clusters were distinct (Silhouette score ∼ 0.55). **D**. To understand the effects of neuromodulation in terms of excitability dynamics, we plotted the `V` (solid) and `w` (dashed) nullclines, with their intersection stable (filled) or unstable (empty) fixed points. With respect to control (orange), MCh (violet) increases the excitability of the neuron by lowering the slope of the *V* -nullcline left branch, while also reducing the adaptation rate (*w*-nullcline slope).

Next, guided by the hypothesis that different neuromodulatory conditions would consistently alter the feature space of a neuron, we submitted the parameter sets to a dimensionality reduction technique – principal component analysis (PCA) – to identify and group the effects of neuromodulators on the AdEx parameters (for example Fig. 1C). The hypothesis implies that in this reduced space, we should observe that parameters for different neuromodulatory conditions cluster into different groups. Indeed, for all considered cases, the parameters characterising the control condition were statistically different from those representing the modulated condition. To further test the validity of this approach, we took advantage of three traces recorded from the same layer III human cortical neuron when stimulating it with the same current protocol (***McCormick and Williamson, 1989***, their Fig.1 A, B, and D). We treated the recordings as three independent traces to fit. Then, we applied the PCA and observed that the groups were overlapping (with Silhouette scores < 0), suggesting that the models were indeed not independent (see Methods and Supplementary Fig. 3).

Then, we analysed how neuromodulation alters neural dynamics by studying how different parameter sets change the system phase plane (Fig. 1D). In phase plane analysis, we plot how the variables *V* and *w* evolve over time and in relation to each other. To understand how neuromodulation influences the behaviour of the system, we study the relationships between these variables when they are slowly changing or remaining steady. To this end, we use the *V* -nullcline (Eq. 3), which represents the set of points where the rate of change of *V* is zero (*dV* /*dt* = 0), and the *w*-nullcline (Eq. 4), which consists of points where *dw*/*dt* = 0.

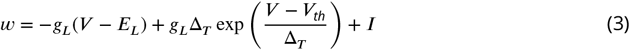

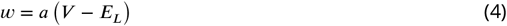

The intersection of the two nullclines defines fixed points – a specific combination of membrane potential (*V*) and adaptation current (*w*) at which both derivatives (*dV* /*dt* and *dw*/*dt*) are zero. The stability of fixed points is given by the eigenvalues of the linearised matrix. If all eigenvalues have negative real parts, the fixed point is stable (attracting). If any eigenvalue has a positive real part, the fixed point is unstable (repelling). If there are both positive and negative eigenvalues, it is a saddle point, and the stability depends on the specific eigenvalues.

In Fig. 1D, we illustrate the insights provided by this approach for a pyramidal neuron in control condition (orange) and under the effect of Methacholine (MCh, violet). In the phase plane, the leak conductance *g*_*L*_ is proportional to the slope of the left branch of the *V* -nullcline (Fig. 1D, solid curves). A steeper slope on the *V* -nullcline and a higher *g*_*L*_ indicate that a small change in the adaptation variable *w* leads to a relatively small change in the membrane potential, resulting in a slower approach to the threshold. Conversely, a shallower slope of the *V* -nullcline implies that a small change in the adaptation variable *w* leads to larger changes in the membrane potential, resulting in a faster approach to the firing threshold. The vertex of the *V* -nullcline marks the point of maximal sensitivity of the membrane potential to the adaptation current *w*. In practical terms, it corresponds to the minimum of the *V* -nullcline, i.e., the lowest voltage at which *dV* /*dt* = 0. The horizontal position of the vertex (in *V*) roughly indicates the threshold region where the neuron is transitioning from rest to spiking. The vertical position (in *w*) reflects the minimum adaptation current needed to keep the neuron at rest at that voltage. Similarly, in the phase plane, the slope of the *w*-nullcline is proportional to the parameter *a* (Fig. 1D, dashed lines). A steeper slope on the *w*-nullcline indicates that a small change in the voltage variable *V* leads to a large change in the adaptation variable, which is a subtracting term in computing the voltage variation *dV*, thus reducing the voltage change and resulting in a slower approach to the threshold. Conversely, a shallower slope of the *w*-nullcline implies that a small change in the adaptation variable *w* results in small changes in the membrane potential, affecting its approach to the firing threshold. The parameter *b* shifts instantaneously *w*, but it does not affect the slope of the *w*-nullcline.

Since MCh is a non-specific cholinergic agonist and mimicks the effect of Acetylcholine (ACh), by comparing the phase planes for control (orange) and modulated (violet) systems (Fig. 1D), we can clearly understand how ACh increases the excitability of the neuron by lowering the *V* -nullcline and *w*-nullcline slopes. Indeed, ACh controls input resistance with receptors opening and/or closing non-voltage-gated *K*^+^ and *Na*^+^ channels. Combinations of these “leak” permeabilities, *g*_*L*_ (in the AdEx), provide a mechanism to generate different excitability states over time (***McCormick, 1989***).

Similarly, voltage-dependent muscarinic currents (IM), *w* (in the AdEx), have a slow component activated upon depolarisation, controlled by the parameter *a* (in the AdEx), which is modulated by ACh (***Benda and Herz, 2003; Kaczmarek et al., 1987***, p.30,165). The magnitude of spike-induced IM, controlled by parameter *b* (in the AdEx), is also modulated by ACh (***Cerina et al., 2015***).

### Cortical pyramidal neurons

We applied our parameter search procedure to the traces from the work of ***McCormick and Williamson, 1989*** on human cortical pyramidal neurons in control conditions (Fig. 1A) and under the bath application of Histamine (HT, Fig. 2A), Methacholine (MCh, Fig. 2B, as Fig. 1B), Norepinephrine (NA, Fig. 2C), and Serotonin (5-HT, Fig. 2D). We extracted the electrophysiological features (as in Fig. 1A and B); fitted AdEx parameter sets for each condition, applied dimensionality reduction to ease their grouping (Fig. 2E), and finally studied their dynamics (Fig. 2F). Overall, compared to the control condition (Fig. 1A), all tested neuromodulators altered spiking frequency in different ways, and influenced other parameters to different extents.

Histamine, HT (Fig. 2A, and 2E dark green points, boundary, and shaded area), is the neuro-modulator that impacts the least firing rate and other features of pyramidal neurons’ firing. From the reduced PCA space and the phase space (Fig. 2E and F) we can see that, with respect to control conditions, HT does not alter the leak conductance (*g*_*L*_) and the spike-triggered adaptation (*b*), but it decreases the slope of the *w*-nullcline (parameter *a*) and the adaptation time constant (*τ*_*w*_). The parameter *a* bundles adaptation conductances responsible for voltage-gated *K*^+^ currents (M-type currents), *Ca*^2+^ currents, and *Ca*^2+^-gated *K*^+^ (AHP-type) currents (***Benda and Herz, 2003***). As such, to capture the complex interactions of these currents, it can also take negative values in the AdEx model — a mathematical abstraction since conductances are not physically negative. HT decreases *Ca*^2+^-activated *K*^+^ conductances, resulting in small depolarization and increased input conductance (***Haas and Konnerth, 1983; McCormick and Williamson, 1991; Flik et al., 2011***).

**Figure 2.**
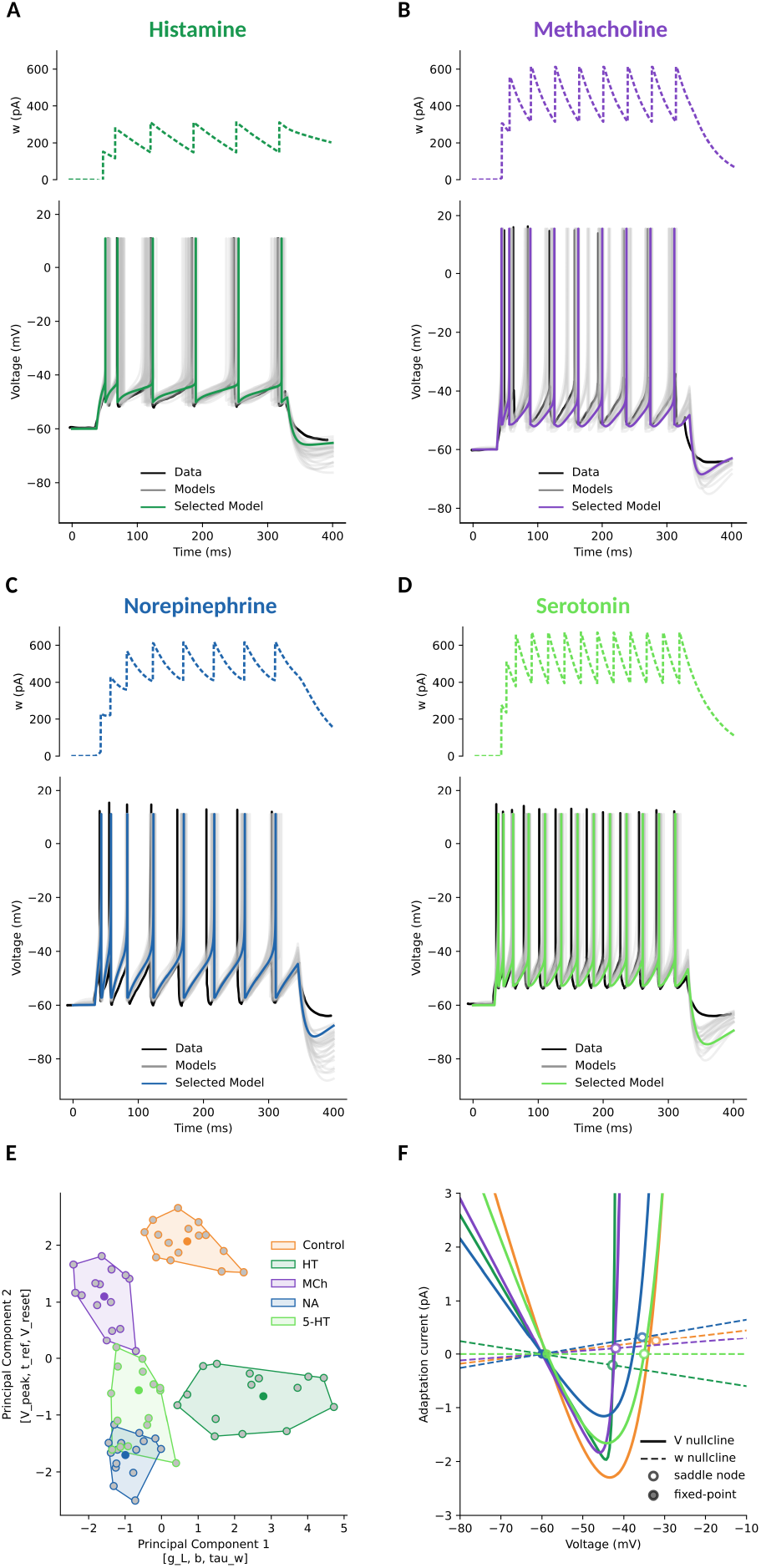
Pyramidal neuron excitability is enhanced by neuromodulation. With respect to the control condition (in Fig. 1A): **A**. Histamine (HT) reduces the inter-spike-interval (ISI). **B**. Methacholine (MCh) reduces more ISI and time-to-first-spike (TFS). **C**. Norepinephrine (NA) has similar effects of reduced ISI and TFS. **D**. Serotonin (5-HT) has the largest impact on both ISI and TFS. In all panels from A to D, top: model adaptation variable *w*, bottom: Voltage, with the original scraped data (black), fitted models (grey), and the selected models (same colours throughout the paper). **E**. To study the relationship between the conditions, we applied a dimensionality reduction technique (PCA) to the parameters of fitted (grey points) and selected (centroid, same colour as in the respective panels) models. In this space, the control (orange) and modulated clusters were distinct (Silhouette score ∼ 0.4). The top three features contributing to the first principal component were [`g_L, b, tau_w`] and the top contributors of the second principal component were [`V_peak, t_ref, V_reset`]. HT cluster presents the largest changes along the first principal component, while 5-HT and MCh have the largest variation along the secondary principal component. **F**. To understand the changes in excitability dynamics, we plotted the *V* (solid) and *w* (dashed) nullclines, with their intersection stable (filled) or unstable (empty) fixed points. With respect to control (orange), all neuromodulators increase the excitability of the neuron by lowering the slope of the *V* -nullcline left branch and raising the vertical position (in *w*) of the *V* -nullcline vertex, which reflects the minimum adaptation current needed to keep the neuron at rest at that voltage. While HT (dark green), MCh (violet), and 5-HT (light green) reduce the adaptation rate (lowering the *w*-nullcline slope), NA (blue) increases adaptation but also further raises the vertex of the *V* -nullcline.

Methacholine, MCh, a non-specific cholinergic agonist (Fig. 2B, and 2E violet points, boundary, and shaded area) has been described above (Fig. 1B). Here we just note the effects of reduction of leak conductance (*g*_*L*_), adaptation decay time (*τ*_*w*_) and its unitary increase (*b*), common to all neuromodulators. As said above, this increases neuron excitability by lowering the *V* -and *w*-nullcline slopes. In contrast to other neuromodulators, MCh does not alter the spike upswing rate (Δ_*T*_) and peak (*V*_*peak*_) with respect to the control condition.

Norepinephrine, NA (Fig. 2C, and 2E blue points, boundary, and shaded area), lowers the leak conductance (*g*_*L*_), the adaptation decay time (*τ*_*w*_) and its unitary increase (*b*). At the same time, it raises the exponential spike raise time (Δ_*T*_). These changes are reflected in the phase space (Fig. 2F). NA increases the neuron’s excitability by lowering the slope of the left branch *V* -nullcline. The shallower *V* -nullcline slope would result in a faster approach to the firing threshold. But, at the same time, NA is the only neuromodulator that raises the slope on the *w*-nullcline. Thus, a small change in the voltage variable *V* leads to a large change in the adaptation variable, subtracting more from the voltage variation *dV*, reducing the voltage change and balancing the effects of a shallow *V* -nullcline. This results in a small relative increase in spiking frequency (control ∼ 13Hz vs. NA 26 Hz). Indeed, as reported by ***McCormick and Williamson, 1989*** and ***McCormick et al., 1991***, the activation of *β*-adrenergic receptors in pyramidal neurons results in an enhanced excitability, mediated by a slow *Ca*^2+^-activated *K*^+^ current.

Serotonin, 5-HT (Fig. 2D, and 2E light green points, boundary, and shaded area), more than MCh but less than NA, lowers the leak conductance (*g*_*L*_), the adaptation decay time (*τ*_*w*_) and its unitary increase (*b*), and raises the exponential spike raise time (Δ_*T*_). In the phase space (Fig. 2F), 5-HT does not alter much the slope of the V-nullcline but increases the excitability of the neuron by flattening the slope on the *w*-nullcline (values of *a* are close to 0). Thus, changes in the voltage variable *V* have little effect on the adaptation variable, which in turn affects the voltage variation *dV* less, resulting in no firing adaptation. Indeed, ***McCormick and Williamson, 1991*** reported that 5-HT suppresses IM currents. At the time, they could not identify the mechanism of suppression. They knew it was not through currents also under control of muscarinic receptors, nor through suppression of *Ca*^2+^ or *Na*^+^-activated *K*^+^ currents. Several recent works (***Amargós-Bosch et al., 2004; De Almeida and Mengod, 2008; Muzerelle et al., 2016***) have shown that 5-HT has an excitatory effect through the activation of 5-HT_*1A*_ receptors.

### Striatal projection neurons

The striatum is the largest nucleus and main input stage of the Basal Ganglia. It receives input from the cerebral cortex, thalamus and midbrain. Around 95% of the neurons in the rodent striatum are striatal projection neurons (SPNs), and the remaining 5% are interneurons (***Graveland and DiFiglia, 1985***). SPNs are approximately equally divided into direct striatal projection neurons (dSPNs) and indirect striatal projection neurons (iSPNs). The dSPNs express D1 dopamine receptors, which have a high density in the striatum and when activated by Dopamine (DA) increase the neuron excitability. On the contrary, the iSPNs express D2 dopamine receptors, which decrease the neuron excitability when activated (***Gerfen and Surmeier, 2011***).

We applied our parameter search procedure to the available experimental data on the effects of DA on SPNs from ***Planert et al., 2013*** (their Fig 4. C, control and after bath application of DA in mice model) to capture the dichotomous effects of DA on AdEx neuronal excitability (Fig. 3). In addition, we applied the same procedure to the experimental data on the effects of Acetylcholine (ACh) from ***Perez-Rosello et al., 2005*** (their Fig. 8 A and B, in rat model).

**Figure 3.**
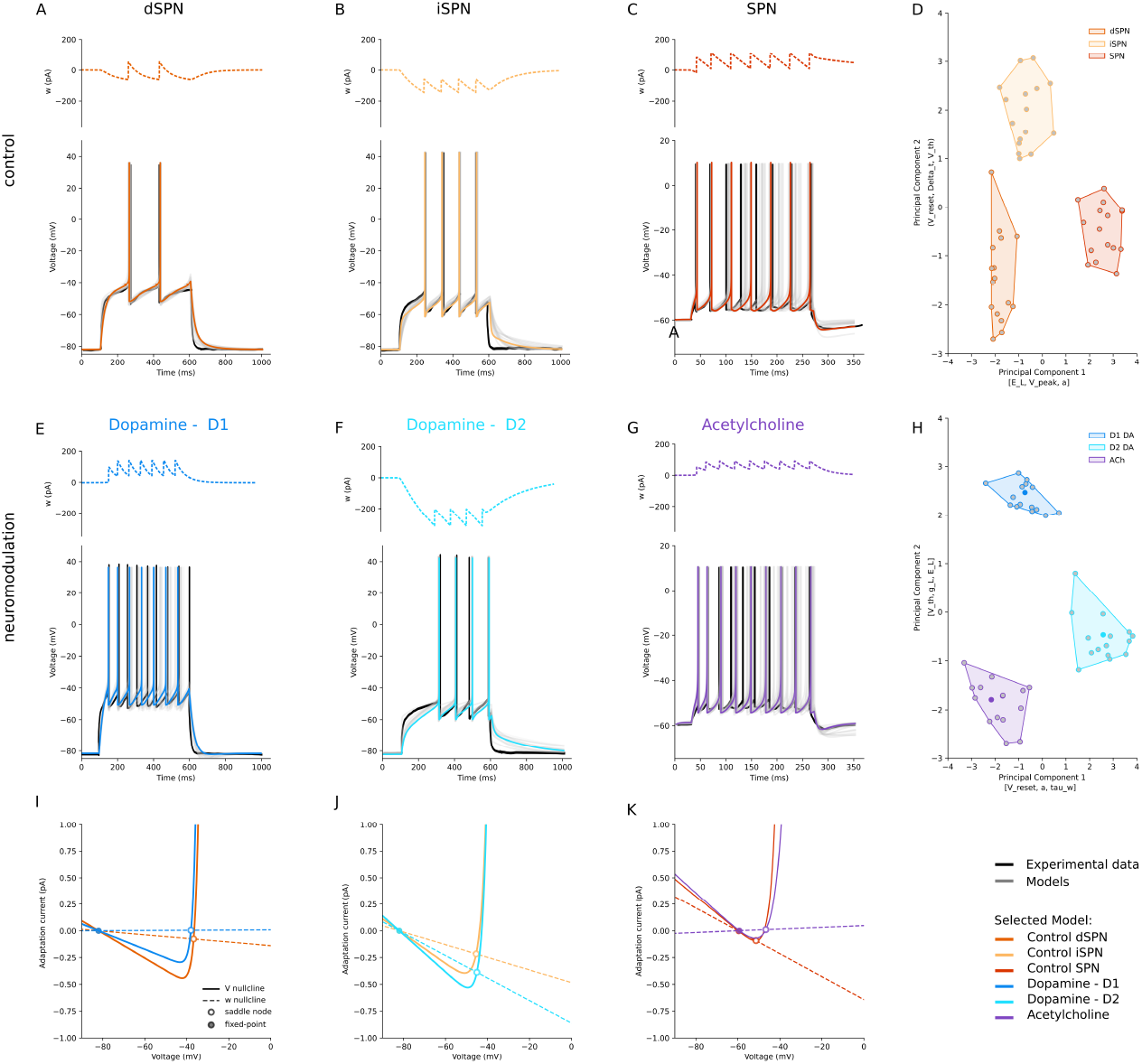
Striatal neurons sharpen and reverse their differences. D1-receptor expressing SPN (dSPN) and D2-receptor expressing SPN (iSPN) behave differently under the effects of Dopamine (DA). In control conditions, dSPNs **(A)** have low, non-adapting spiking rates, with long latency to spike, and iSPNs **(B)** have a similar behaviour **C**. Regardless of their receptor expression, more depolarised conditions only increase the firing rate of SPNs without increasing spiking adaptation. **D**. In the PCA-reduced parameter space, the behaviour of dSPN (dark orange) and iSPN (light orange) is distinguished by approach to threshold and after-spike reset. A different holding voltage, without considering dopamine receptor expression (red), shifts the parameters along the Principal Component 1 axes [`E_L, V_peak, a`]. The Silhouette score is ∼ 0.68. **E**. In DA bath conditions, D1-expressing SPNs lose the latency to first spike and triple their spiking rate. **F**. Conversely, in the same condition, D2-expressing SPNs increase their latency and not their firing. **G**. Acetylcholine (ACh) increases the firing rate of SPNs and introduces spike adaptation. **H**. In the PCA space, DA increases the difference between dSPNs and iSPNs (Silhouette score in control condition is ∼ 0.59, while under the effect of DA is ∼ 0.66) and in general improves the grouping of the clusters (Silhouette score goes from ∼ 0.68 to ∼ 0.74). **I-K**. In the phase space, it is clearly visible the opposite effects of DA on dSPNs and iSPNs. On dSPNs **(I)**, DA flattens both the left branch of *V* -nullcline and the slope of *w*-nullcline, reducing latency and increasing firing. On iSPNs **(J)**, DA raises the left branch of *V* -nullcline and increases the slope of *w*-nullcline, which results in longer latency. In line with other descriptions, ACh **(K)** increases the excitability of SPNs by lowering the slope of the V-nullcline left branch and flattening the adaptation rate (*w*-nullcline slope).

**Figure 4.**
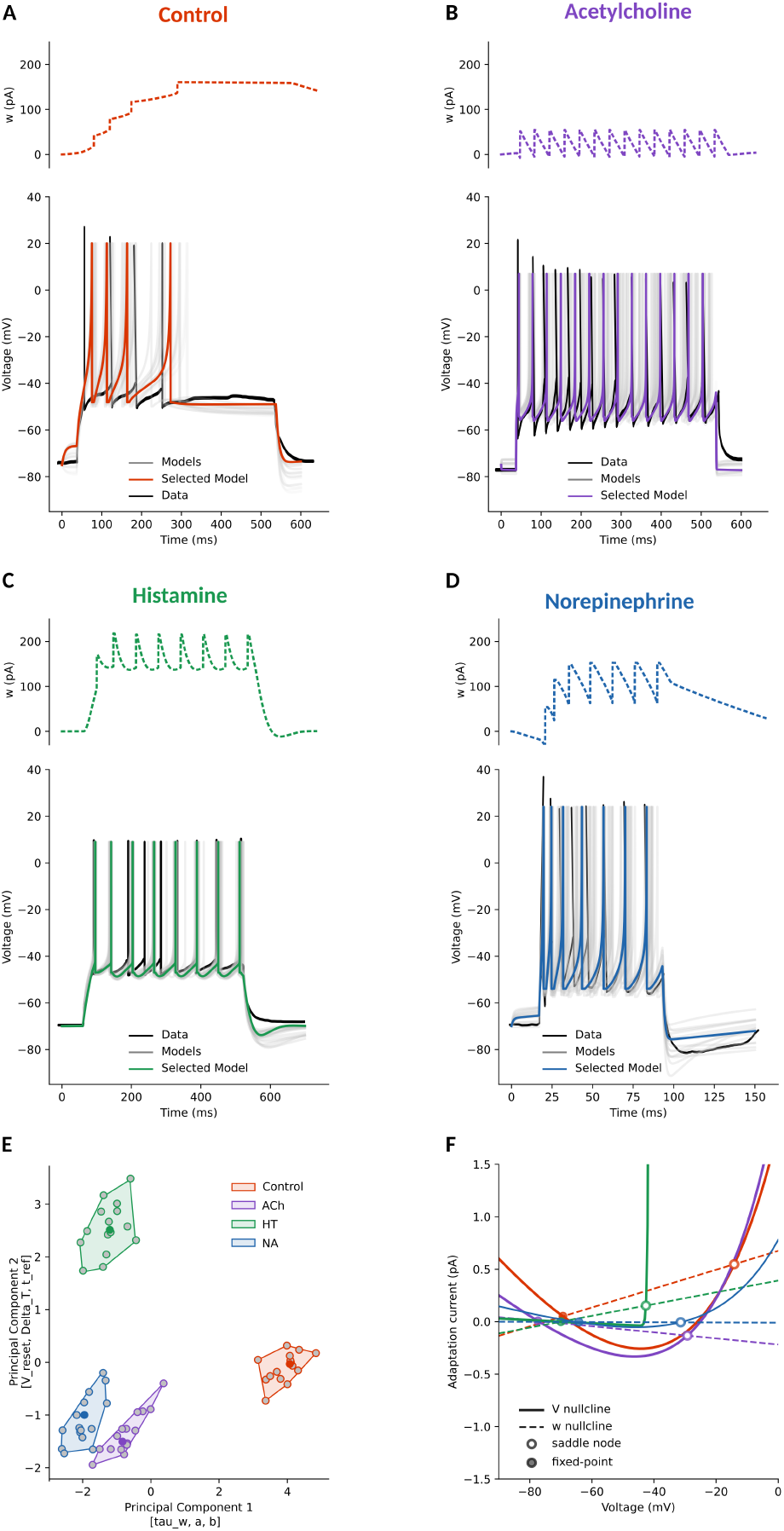
Dentate granule cells (DGCs) have polarised responses to neuromodulators. **A**. In control conditions, DGCs strongly adapt their firing. **B**. Acetylcholine (ACh) increasingly reduces inter-spike-interval (ISI) and abolishes spike adaptation. **C**. Histamine (HT) reduces spike adaptation but leaves ISI close to control conditions. **D**. Norepinephrine (NA) has effects of reduced spike adaptation and ISI similar to ACh. **E**. In the PCA-reduced parameter space, the control (red) and modulated clusters were distinct (Silhouette score ∼ 0.6). HT cluster (light green) presents the largest variation along both Principal Component axes, and NA and ACh have similar variations. The parameters most influencing PC1 are related to subthreshold and adaptation properties [`Delta_T, b, tau_w`], while PC2 is primarily shaped by spiking threshold and reset dynamics [`V_th, V_peak, V_reset`]. **F**. In the phase space, we have the *V* (solid) and *w* (dashed) nullclines, with their intersection stable (filled) or unstable (empty) fixed points. With respect to control (red), all neuromodulators, except ACh, increase the excitability of the neuron by lowering the slope of the *V* -nullcline left branch and raising the vertical position (in *w*) of the *V* -nullcline vertex. While HT (dark green) and NA (blue) reduce the adaptation rate (lowering the w-nullcline slope), ACh increasingly removes adaptation (captured by the negative *w* slope) and also lowers the vertex of the *V* -nullcline.

We grouped the six models into two conditions: control (Fig. 3A-C, different shades of orange for dSPN, iSPN, and SPN) and under the influence of DA and ACh (Fig. 3E-F, DA-molulated dSPN in blue, DA-modulated iSPN in cyan, and Fig. 3G, ACh-modulated SPN in violet), and we performed PCA on the control and modulated parameter sets, as shown in Fig. 3D and H, respectively. In both cases, the models form distinct clusters, but neuromodulation increases the separation between them, as indicated by a rise in the Silhouette score from ∼ 0.68 to ∼ 0.74. Finally, we investigated the effect of DA and ACh on the corresponding control models by examining their dynamics in the phase plane (Fig. 3 I-K).

In dSPNs (Fig. 3E), DA has an excitatory effect. It decreases the leak conductance *g*_*L*_, resulting in a shallower slope of the *V* -nullcline, implying that a small change in the adaptation variable *w* leads to larger changes in the membrane potential, resulting in a faster approach to the firing threshold. It also makes the slope of the *w*-nullcline shallower, such that a change in the adaptation variable *w* leads to smaller changes in the membrane potential, affecting less its approach to the firing threshold. In addition, a decrease in the spike threshold (*V*_*th*_) alongside a reduction in the adaptation parameters *b* and *τ*_*w*_, results in a slower approach to threshold. Indeed, DA enhances *Na*^+^ conductance, acting on *g*_*L*_ (***Planert et al., 2013; Gorelova and Yang, 2000***).

In iSPNs (Fig. 3F), DA has an inhibitory effect. It increases the leak conductance *g*_*L*_, reflecting an increase in *K*^+^ conductance, steepening the slope of the left branch of the *V* -nullcline, resulting in a slower approach to the threshold. It also steepens the slope on the *w*-nullcline such that a small change in the voltage variable *V* leads to a large change in the adaptation variable, which is a subtracting term in computing the voltage variation *dV*, thus reducing the voltage change and resulting in a slower approach to threshold. Moreover, the increase in the adaptation parameters *b* and *τ*_*w*_ and decrease of *a* underscores a heightened delay in the first spike and adaptation response.

On SPNs, without considering the expression of DA receptors, ACh increases excitability compared to control (Fig. 3C and G). In line with the expected influence of ACh, the AdEx models exhibit a decrease in the leak conductance *g*_*L*_, representative of enhanced *Na*^+^ channel permeability, and *V*_*th*_, indicative of a facilitated response to incoming stimuli.

In summary, DA can modulate *Na*^+^, *K*^+^, and *Ca*^2+^ channels. An increase or decrease in *g*_*L*_ might reflect an increase or decrease in *Na*^+^ and *K*^+^ conductance, leading to hyperpolarization or depolarization. Changes in the adaptation parameters (*a, b*, and *τ*_*w*_) could suggest alterations in *Ca*^2+^ channel activity affecting neuron’s firing patterns. It is important to stress that the parameters *g*_*L*_, *a, b, τ*_*w*_, *V*_*reset*_ are modulated in the opposite directions depending on the DA receptor type (See Supplementary Material 2).

### Hippocampal neurons

In the hippocampal formation, the Dentate Gyrus (DG) occupies an important input position, receiving from the entorhinal cortex and projecting to the first portion of the Cornus Ammonis (CA1). The principal neurons of the DG are the granule cells.

To characterize dentate granule cells, we applied our parameter search procedure to electro-physiological traces from the works of ***Carver and Shapiro, 2019*** (their Fig. 5, control vs bath application of oxo-M, a muscarinic agonist, mimicking ACh conditions, in mice), ***Greene and Haas, 1990*** (their Fig. 3, HT condition, in rats), and ***Haas and Rose, 1987*** (their Fig. 2, bath application of Isoproterenol, a *β*-adrenergic agonist, mimicking NA condition, in rats). We report the fitting for the control condition (Fig, 4A) and under modulation of ACh (Fig. 4B), HT (Fig. 4C), NA (Fig. 4D). As before, we measured the electrophysiological features; we obtained AdEx parameter sets for each condition through fitting, and then we applied dimensionality reduction to ease their grouping (Fig. 4E). Once we found these groups, we studied their dynamics (Fig. 4F).

**Figure 5.**
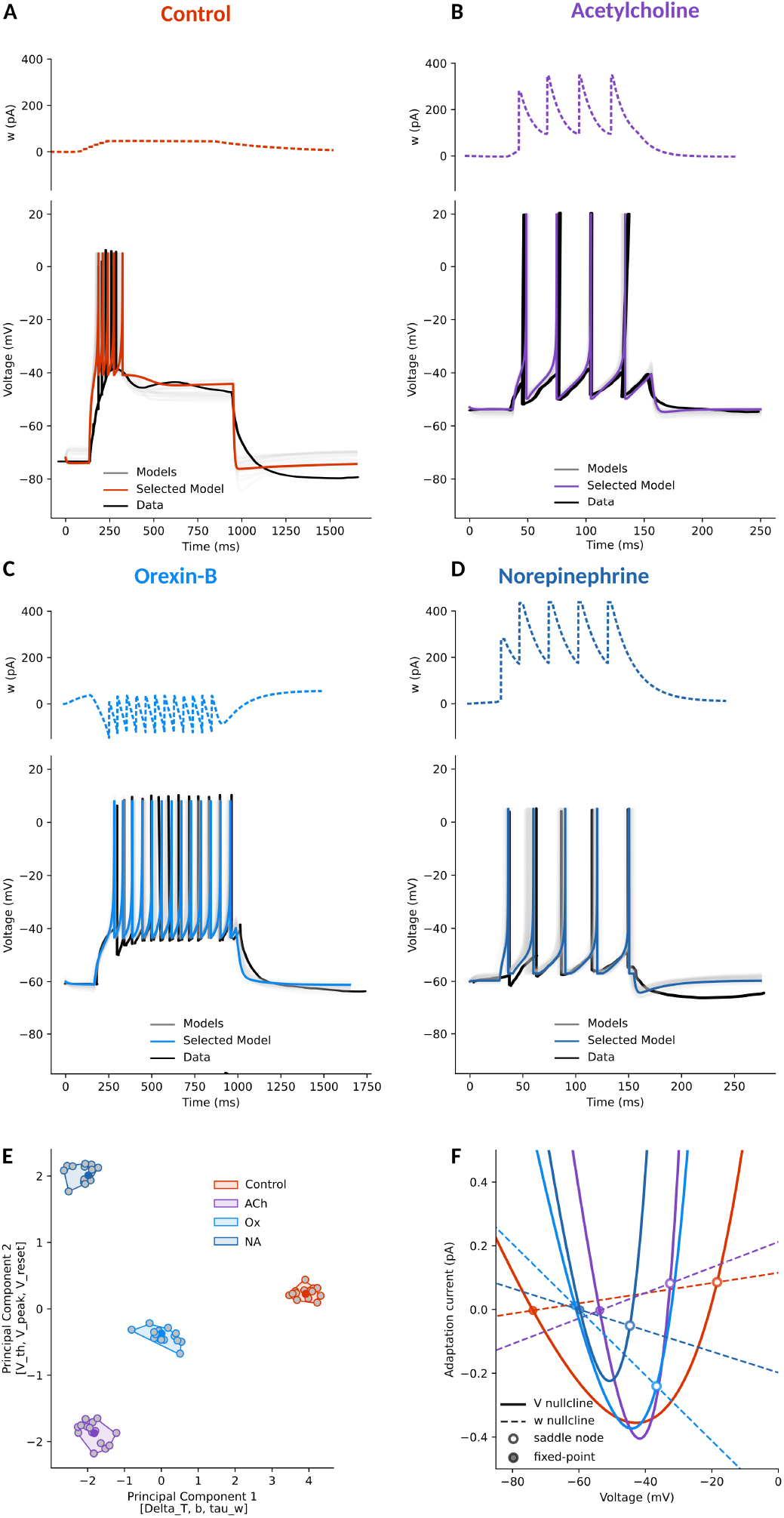
The switch between bursting and tonic firing of Thalamocortical Projecting Neurons (TPS). **A**. In control conditions, TPNs strongly adapt their firing. A spiking behaviour known as “bursting”. All neuro-modulators make the TPNs switch to a regular spiking behaviour known as “tonic” firing. **B**. Acetylcholine (ACh) increases the inter-spike-interval (ISI) and abolishes spike adaptation. **C**. Orexin-B (Ox) too abolishes spike adaptation and introduces latency for the time to first spike. **D**. Norepinephrine (NA) also reduces spike adaptation and has ISI similar to ACh. **E**. In the PCA-reduced parameter space, the control (red) and modulated clusters were clearly separated (Silhouette score ∼ 0.86). NA and ACh have similar variation along Principal Component 1 (whose top influencing parameters are [`Delta_T, b, tau_w`] but differ along the Principal Component 2 (which is mainly influenced by [`V_th, V_peak, V_reset`]). **F**. In the phase space, we can see that TPNs’ response to all neuromodulators is a narrower *V* -nullcline (solid). This corresponds to a slower approach to threshold, a less excitable system, notwithstanding the raised *E*_*L*_ (a known effect of these neuromodulators on TPNs). Indeed, note that in the different temporal scales, all firing rates are roughly equal (∼ 4 sp/s). The slopes of the *w*-nullcline for the different conditions explain various spiking behaviours. In control (red), the positive *w* slope brings strong adaptation (“bursting”). In ACh condition (violet), the even steeper *w*-nullcline is balanced by the narrower *V* -nullcline, resulting in a slower approach to threshold, which allows the decay of *w* and the absence of adaptation (“tonic”). In Ox conditions (light blue), the (relatively strong) negative *w*-nullcline captures the latency and absence of adaptation. In NA conditions (dark blue), the negative *w*-nullcline balances the higher *V* -nullcline vertex, which allows higher adaptation currents (panel D, top).

In the original work from ***Carver and Shapiro, 2019***, excitatory input currents of 60, 100, 140 pA were used in both the control and ACh conditions. We fitted our model to all the values in the control condition, and the 140 pA data because of its higher quality and consistency. The authors also reported the use of a holding current. However, they did not specify its intensity. For our fitting, we have assumed −300 pA for the control and −200 pA in the condition of muscarinic agonist oxo-M bath application. This is because ACh increases the input resistance, and less current is usually needed to hold the neurons at the desired voltage (−75 mV, here and in the original article).

In control conditions (Fig. 4A, orange point and shaded area in E, and red lines in F), dentate granule cells strongly adapt their firing. In this condition, the left branch of the *V* -nullcline is high, with high leak currents, and the *w*-nullcline has a high positive slope, providing a powerful adaptation current. Our fitting also places high values for the *τ*_*w*_, stretching the effects of even low *b* unitary increases to *w*.

Compared to the control condition, the bath application of muscarinic agonist oxo-M (Fig. 4B, violet points and shaded area in E, and violet lines in F) increases the excitability of dentate granule cells by lowering the left branch of the *V* -nullcline, and bringing the *w*-nullcline to negative slope (*a*). The effects of ACh are similar on granule cells and cortical pyramidal neurons. Indeed, as for pyramidal neurons, ACh controls non-voltage-gated *K*^+^ and *Na*^+^ channels (***Carver and Shapiro, 2019; Brown and Passmore, 2009***), and it also slows voltage-dependent muscarinic-dependent *Ca*^2+^-activated adaptation currents (***Carver and Shapiro, 2019; Vogt and Regehr, 2001***).

HT application (Fig. 4C, green points in E, and green lines in F) does not lead granule cells to an increase in spiking frequency as strong as ACh and NA. However, it shifts the parameter space of the dentate granule cells in a different direction. It lowers the left branch of the *V* -nullcline, and therefore its leak currents. But the *w*-nullcline has a positive slope close to the control condition, which provides a powerful adaptation current, balanced by low values for *τ*_*w*_, which shortens the effects of *b* unitary increase to *w*. At the same time, HT raises the reset voltage (*V*_*reset*_) and the spike upswing rate (Δ_*T*_), leading to smaller depolarisation and a slower approach to threshold. Indeed, ***Greene and Haas, 1990*** showed that HT causes, through specific H_2_-receptors, a block of the long-lasting component of spike after-hyperpolarisation (AHP) and a reduction of firing adaptation. In addition, HT selectively blocks the late long-lasting *Ca*^2+^-dependent outward tail current without any reduction of inward current, as captured by the model *V* - and *w*-nullclines.

NA has a similar effect to ACh on dentate granule cells (Fig. 4D, blue point and shaded area in E, blue lines in F). NA increases the excitability of dentate granule cells by lowering even more than ACh the left branch of the *V* -nullcline, and bringing the *w*-nullcline to 0 (or negative) slopes. The similarity between the two neuromodulators is clear in Fig. 4E, where the two parameter clusters (violet for ACh and blue for NA) are distinct from one another but close compared to control and HT conditions. Indeed, ***Haas and Rose, 1987*** showed that NA reduces dentate granule cell spike adaptation through *β*-receptors blockade of *Ca*^2+^-dependent *K*^+^ conductance.

### Thalamocortical projecting neurons

The thalamus is a central structure located in the forebrain above the midbrain. It acts as a central hub, with nerve fibres — the thalamocortical radiation — connecting it to all parts of the cerebral cortex. Nearly all thalamic neurons (except those in the reticular nucleus) project to the cortex, and every cortical region sends feedback to the thalamus, but collateral connections reach the basal ganglia, hippocampus, colliculus, and cerebellum. It relays and processes sensory and motor signals and plays key roles in consciousness, sleep, and alertness (***Jones, 2012***).

To characterise thalamocortical projecting neurons (TPNs), we applied our parameter search procedure to electrophysiological traces from several works. In the absence of neuromodulators, the TPNs of all nuclei display a bursting activity mode. For our fitting of this mode, we used mouse Centro-Lateral neuron data from the work of ***Govindaiah and Cox, 2006***, for their clarity, but analogous voltage responses can be seen in the works of McCormick (***McCormick and Prince, 1987; McCormick, 1989; McCormick and Feeser, 1990***). In all thalamic nuclei, ACh provided by brainstem nuclei (***Moruzzi and Magoun, 1949; Steriade et al., 1990***) causes the switch to tonic firing mode (***Avanzini et al., 2000***). For our fitting, we used the traces from ***McCormick, 1989*** (their Fig. 4A, right). Similar effects are induced by Orexin-B (Ox), a drug targeting channels also targeted by Dopamine, and NA. We used ***Govindaiah and Cox, 2006*** for Ox (their Fig. 5, control vs Orexin-B bath conditions) and ***McCormick, 1989*** for NA (their Fig. 2B).

In our Fig. 5, we report the fitting for the control (5A) and ACh (5B), Orexin-B (5C), NA (5D) conditions. As before, we measured the electrophysiological features, we obtained AdEx parameter sets for each condition through fitting, and then we applied dimensionality reduction to ease their grouping (Fig. 5E). Once we found these groups, we studied their dynamics (Fig. 5F).

In the control condition, where the *E*_*L*_ is low (around −70 mV, Fig. 5A, red points and shaded area in E, and red lines in F), TPNs strongly adapt their firing. The low *E*_*L*_ coupled with relatively high *g*_*L*_ and adaptation *a* all concur in a slow approach to threshold. The left branch of the *V* -nullcline is high, with high leak currents, and the *w*-nullcline has a relatively high positive slope, providing a powerful adaptation current. The absence of ACh in the thalamus reduces TPNs excitability due to a muscarinic receptor-mediated increase in non-voltage dependent *K*^+^ conductance that lowers the resting membrane potential (***McCormick, 1989***). When stimulated in this condition, TPNs generate a *Ca*^2+^ spike that triggers a burst of fast *Na*^+^/*K*^+^ action potentials (***McCormick, 1989***). The bath application of ACh (Fig. 5B, violet points and shaded area in E, and violet lines in F) reduces the adaptation of TPNs and increases their excitability due to the reverse mechanism described above, muscarinic receptor-mediated decrease in *K*^+^ conductance (***McCormick, 1989***). In these conditions, TPNs fire tonically, with ACh depolarising the neuron out of the voltage range in which the low threshold *Ca*^2+^ current is active (Fig. 4A in ***McCormick, 1989***). The AdEx model does not contain a term explicitly representing *Ca*^2+^ currents, but the sub-threshold adaptation *a*, the low *E*_*L*_ coupled with relatively high *g*_*L*_, and non-zero adaptation *a* all concur in a slow approach to threshold. In the model, this is captured by a narrow V-nullcline.

A different picture holds for the other neuromodulators, while resulting in a similar tonic firing mode. The application of Orexin-B (Fig. 5C, data trace in black, light blue points, shaded area in E, and lines in F) is characterised by latency to the first spike and regular (“tonic”) firing. This is due to a decrease in the leak potassium current (***Govindaiah and Cox, 2006***). The AdEx model does not contain a term representing the influence of Orexin-B on calcium currents, but the relatively low *g*_*L*_ and negative sub-threshold adaptation *a* concur in the first slow approach to threshold. After the first spike, when the *b* term is added to *w*, the narrow *V* -nullcline balances the adaptation term by slowing the dynamics and allowing the decay of *w*, contributing to the tonic firing mode.

Also the application of NA (Fig. 5D, data trace in black, dark blue points, shaded area in E, and lines in F) shifts the TPNs from the bursting to tonic mode of activity. As described by ***McCormick, 1989***, this is due to a *α*_1_-adrenoceptor-coupled decrease in *K*^+^ conductance (their Fig. 2 B and C). This decrease induces a slow depolarisation bringing the membrane potential closer to the firing threshold, increasing the input resistance, and time constants. In the AdEx model, the narrow *V* -nullcline balances the adaptation term by slowing the dynamics and allowing the decay of *w*, effectively mimicking the increase in input resistance and time constants, thus contributing to the tonic firing mode.

### Thalamic reticular neurons

The thalamic reticular nucleus encircles the thalamus laterally and dorsally. Reticular cells are GABAergic and integrate cortical (***Bourassa and Deschênes, 1995***), brainstem and mesencephalic inputs (***Spreafico et al., 1993***). Reticular cells project outputs directly, and exclusively, to the thalamus (***McAlonan et al., 2006; Wang et al., 2001***).

To characterise thalamic reticular neurons (TRNs), we applied our parameter search procedure to electrophysiological traces from the works of ***McCormick and Prince, 1986***. However, they applied orthodromic synaptic stimuli to the neurons instead of current injections (used in all other datasets and in our modelling framework). This causes a slower decay of the membrane, which is the effect we see when comparing the models to the data (Fig. 6A and B).

**Figure 6.**
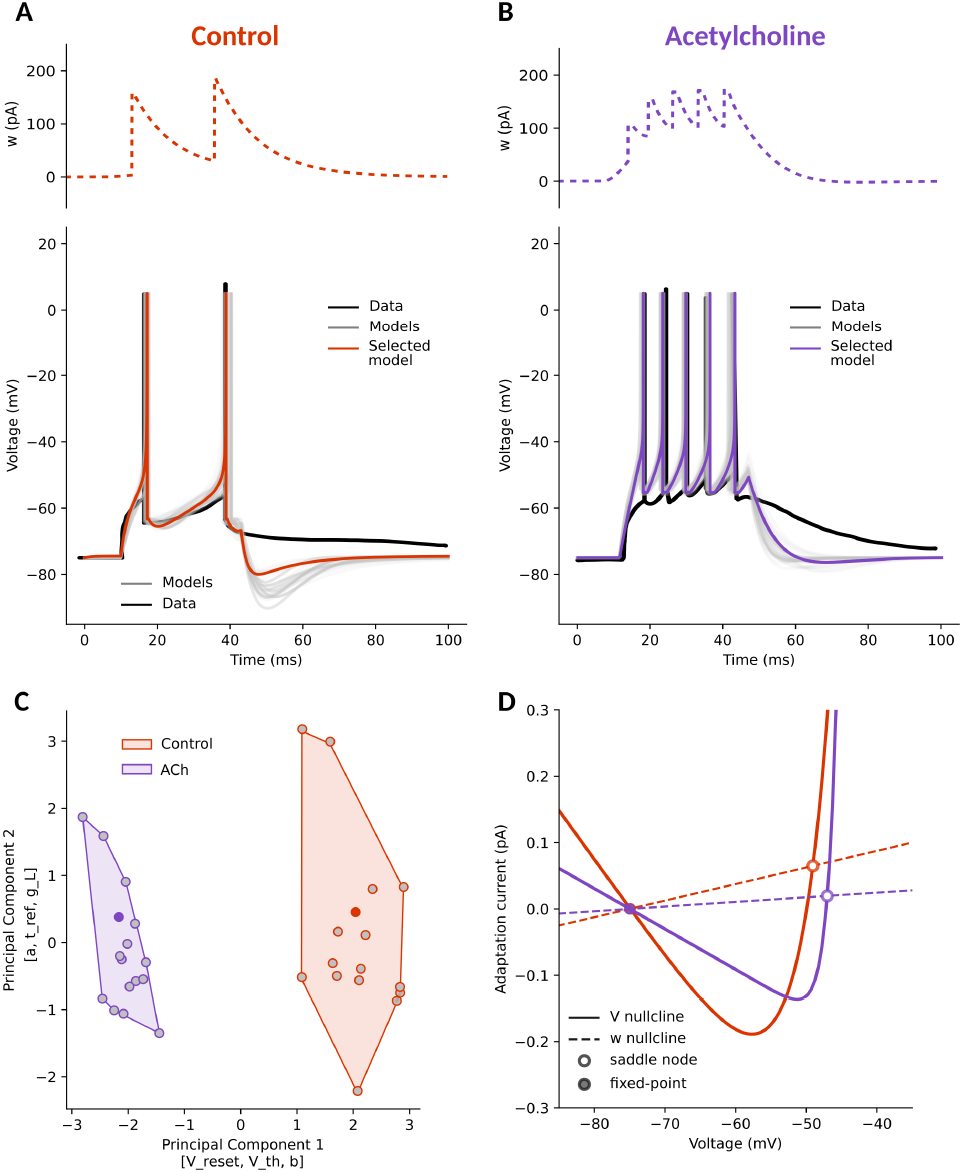
The switch between bursting and tonic firing of Thalamic Reticular Neurons (TRNs). **A**. In control conditions, a TRN strongly adapts its firing. **B**. The same TRN, in Acetylcholine (ACh) conditions, does not adapt its firing, raises the resting potential, and has a shorter time to first spike latency. **C**. In the PCA-reduced space, the control (red) and ACh-modulated (violet) clusters were distinct (Silhouette score ∼ 0.67), mainly shifting along the first component [`V_reset, V_th, b`]. **D**. In the phase space, with respect to control, ACh increases the excitability of the neuron by lowering the slope of the left branch *V* -nullcline, reducing the slope of *w*-nullcline, while also increasing the threshold zone by shifting the *V* -nullcline vertex to more negative values.

In control conditions (Fig. 6A, orange point and shaded area in C, and red lines in D), reticulate neurons adapt their firing. In this condition, the left branch of the *V* -nullcline is high, with high leak currents, and the *w*-nullcline has a high positive slope, providing a powerful adaptation current. From ***McCormick and Prince, 1986*** we know that the absence of ACh in the reticular thalamus reduces reticular neurons excitability due to a low muscarinic receptor-mediated *K*^+^ conductance. In these conditions, thalamic reticular neurons fire in single spike mode (***McCormick and Prince, 1986***, their Fig. 3A).

Compared to the control condition, the bath application of ACh in the reticular thalamus (Fig. 6B, violet points and shaded area in C, and violet lines in D) causes a membrane depolarization by lowering the left branch of the *V* -nullcline, and reduces spike adaptation by also lowering the slope of *w*-nullcline. The effects of ACh are presumably mediated through an increase in *K*^+^ conductance (***McCormick and Prince, 1986***, their Fig. 3A).

Considering together the thalamic relay and reticular neurons results, an interesting network property emerges. In the absence of ACh modulation, thalamocortical neurons exhibit brief spike bursts in response to stimuli. This is matched with the relatively modest single-spike responses from reticular neurons, potentially enabling stimuli to propagate further within the network. With the introduction of ACh, thalamocortical neurons show heightened tonic single-spike firing in response to stimuli. This augmented firing is mirrored by a corresponding increase in firing among reticular neurons. These findings tentatively propose that while neuromodulation reshapes the intrinsic properties and behaviours of individual cells, it may also trigger compensatory adaptations within the broader network.

### Cerebellar neurons

In the cerebellar cortex, Purkinje cells (PCs) are GABA-ergic neurons, essential for controlling motor activity by directly inhibiting deep cerebellar nuclei projecting to brainstem premotor areas (***Walter and Khodakhah, 2006; Manto et al., 2012; Judd et al., 2021***).

To characterize PCs, we applied our parameter search procedure to electrophysiological traces from the works of ***Fleming and Hull, 2019*** for control and 5-HT conditions (their Fig. 1F); and ***Watkins and Mathie, 1996*** for ACh condition (their Fig. 6C). In the latter, Muscarine was bath applied, however, since Muscarine is a selective agonist of ACh it provides a good approximation of the modulatory effects mediated by ACh receptors. We report the fitting for the control condition (Fig, 7A) and under the bath application of Muscarine (Fig. 7B), Serotonin (Fig. 7C). As before, we measured the electrophysiological features; we obtained AdEx parameter sets for each condition through fitting, and then we applied dimensionality reduction to ease their grouping (Fig. 7D). Once we found these groups, we studied their dynamics (Fig. 7E).

**Figure 7.**
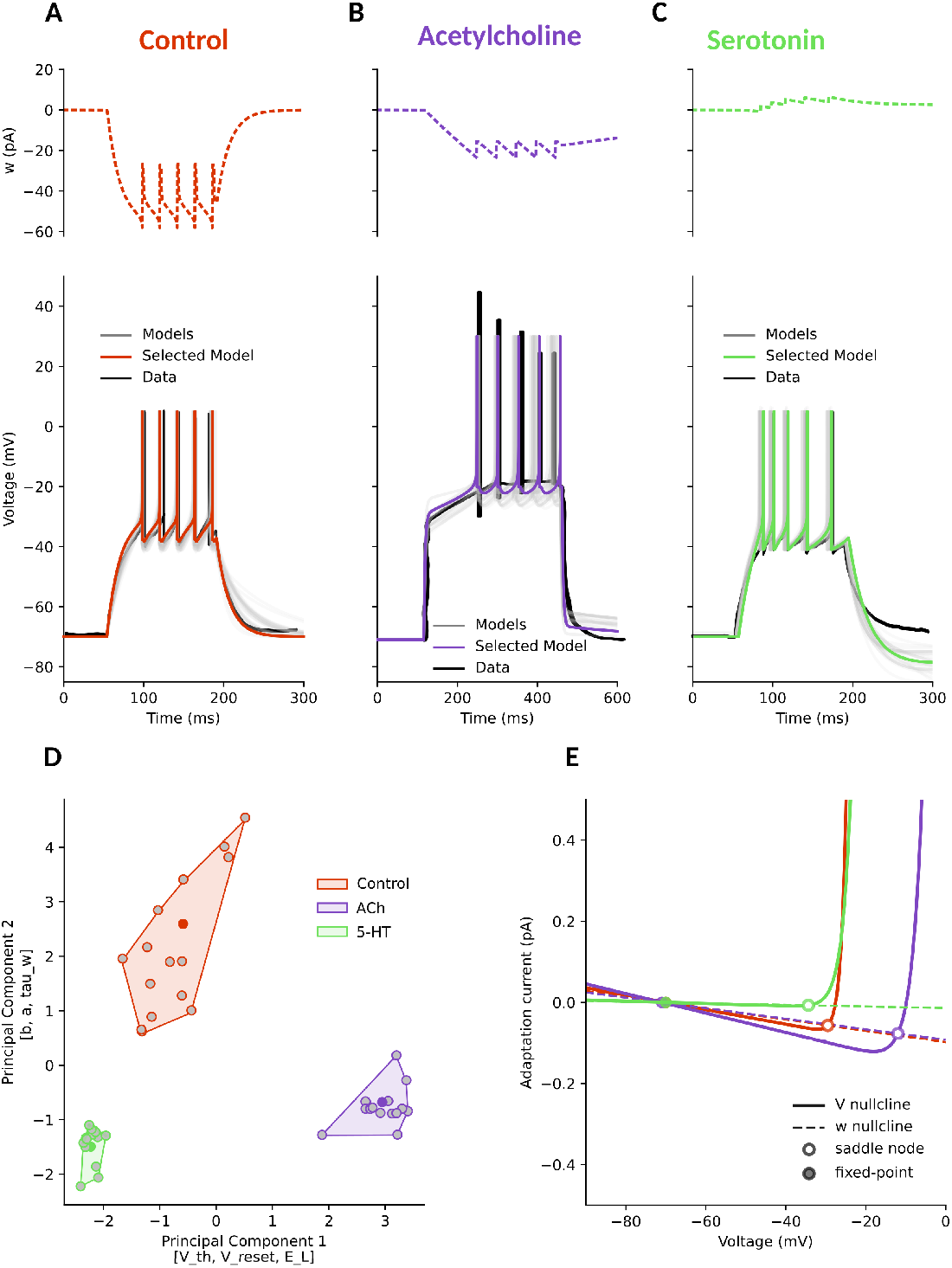
Neuromodulators change the responsiveness of Cerebellar Purkinje cells (PCs). **A**. In control conditions, PCs have a delayed time to first spike (TFS, ∼ 50 ms) and do not adapt. **B**. In Acetylcholine (ACh) conditions, the TFS doubles (∼ 100 ms), and the steady potential rises by ∼ 20 mV. **C**. Serotonin (5-HT) conditions are similar to control conditions, with decreased TFS and increased adaptation. **D**. In the PCA-reduced space, with respect to the control (red) cluster, both modulator clusters were shifted to lower values over the first Principal Component [`V_th, V_reset, E_L`], and the second Principal Component [`b, a, tau_w`]: ACh (violet) on the right, and 5-HT (light green) on the left. They were all distinct (Silhouette score ∼ 0.77). **E**. In the phase space, with respect to control, ACh raises the left branch of the *V* -nullcline, slowing the approach to threshold, and shifts its vertex towards more depolarised values, explaining the raised steady state. 5-HT (light green) brings the *w*-nullcline to positive values and lowers the threshold zone by shifting the *V* -nullcline vertex to more negative values.

**Figure 8.**
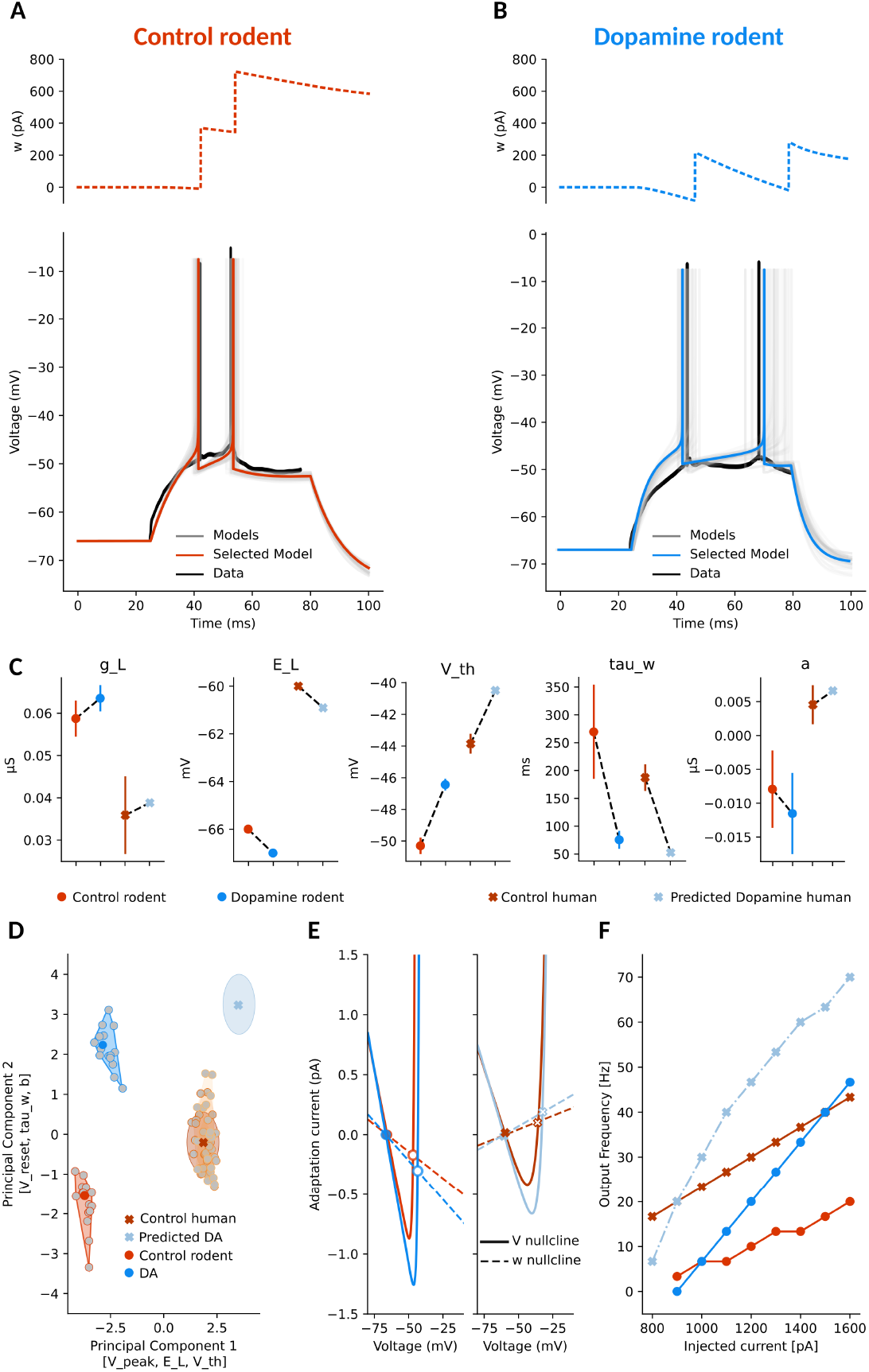
Predicting dopamine effects on humans. AdEx model fitting of voltage traces recorded from rodent cortical neurons in control **A**. and after bath application of DA **B**. The top panels show the evolution of the adaptation current *w*, while the bottom panels show the model voltage traces compared to experimental data. Specifically, thin grey lines represent the pool of fitted models, colored lines the selected best model and black traces are the recorded data. **C**. Comparison of selected parameters across conditions and species. In red and blue circles, rodent control and DA average values are plotted, in dark red crosses the average of human control data, and in light blue crosses the predicted human parameters obtained by applying rodent DA-induced parameter shifts. **D**. PCA of model parameters across conditions. The first Principal Component is mostly influenced by [`V_peak, E_L, V_th`] while the second Principal Component axes by [`V_reset, tau_w, b`]. Controls and DA models form distinct clusters, with DA mainly shifting them along the y-axis. **E**. Phase-plane analysis of the selected AdEx models in rodent (left) and human (right). With respect to control (orange), DA increases the magnitude of the slope of adaptation. **F**. Firing-frequency curve (firing rate vs. input current) for rodent and human models in control and DA conditions.

Purkinje cells, with their extended dendritic tree, have a high capacitance, which characterizes their response to current injection. This is represented in our AdEx models by a negative adaptation current in all conditions, which serves the purpose of slowing the changes in membrane potential. In control conditions (Fig. 7A, orange point and shaded area in D, and lines in E), Purkinje cells present no spiking adaptation. In this condition, the left branch of the *V* -nullcline is shallow, with low leak currents, and the *w*-nullcline has a negative slope, providing little adaptation current.

Compared to the control condition, the bath application of ACh (Fig. 7B, violet points and shaded area in D, and violet lines in E) causes a membrane depolarisation by lowering the left branch of the *V* -nullcline, and shifts the *V* -nullcline vertex towards more depolarised values. In addition, it causes a progressive decrease in the interspike intervals. The effects of ACh are presumably mediated through an increase in *K*^+^ conductance and channel specificity ***Watkins and Mathie, 1996***.

5-HT has an opposite effect to ACh on Purkinje cells (Fig. 7C, green point and shaded area in D, green lines in E). It increases the excitability by lowering the left branch of the *V* -nullcline, and flattening the *w*-nullcline. Moreover, it also shifts the *V* -nullcline vertex to more hyperpolarised values, producing a spiking adaptation effect.

### Predicting the effect of Dopamine on Human Cortical pyramidal neurons

The seminal work by ***McCormick and Williamson, 1989*** did not explore the effect of Dopamine (DA) on human cortical neurons. We therefore took the opportunity to predict it. Specifically, we used data from ***Onn et al., 2006*** (their Fig. 3a1 and a2), which report the effect of DA on rodent cortical neurons, mapped those changes onto the AdEx parameters, and used them to estimate the potential impact of DA on human cortical neurons.

We fitted the AdEx model parameters to match control and DA conditions (Fig. 8 A and B, respectively). Five selected parameters, and how they change under dopaminergic modulation, are illustrated in Fig. 8C. The predicted human DA parameters are illustrated in light blue crosses and are obtained by applying rodent DA-induced parameter shifts to the average human control model (Supplementary Material 2). While we acknowledge the possibility of species-specific differences, we found that the firing frequency curves under control conditions in both rodents and humans exhibit similar slopes (Fig. 8F, orange curves). This similarity suggests that the baseline excitability properties are comparable, supporting our decision to apply the DA-induced parameter changes observed in rodents to the human models.

Based on this approach, we predict that also in humans DA would increase *g*_*L*_, *V*_*th*_, and *V*_*reset*_ and decrease *E*_*L*_, *b, τ*_*w*_, and Δ_*T*_ . The parameter *a*, which controls the subthreshold adaptation, displays opposite baseline values across species. In rodent control models, *a* has a negative average value, and it further decreases under dopaminergic conditions. In contrast, the human control model exhibits a positive average value. This difference opens two possibilities when predicting the dopaminergic effect. We can take into account the increase in the absolute value of *a* or a signed decrease (see Methods for detail and Supplementary Fig. 4). Both approaches produce comparable results in terms of predicted dynamics, while the increase in firing frequency is more pronounced in the second case. In the main text (Fig. 8), we report the simulations using the increase in *a* (absolute value strategy) while in the Supplementary Fig. 4 we also report the case with a decrease in *a* (signed value strategy).

## Discussion

In this study, we demonstrated that neuromodulators reshape neuronal excitability by altering the dynamical parameters of a conductance-based model, the Adaptive Exponential Integrate-and-Fire (AdEx). By extracting core electrophysiological features from published traces and fitting AdEx parameters across multiple neuron types, brain regions, and neuromodulatory conditions, we systematically showed that these changes form distinct clusters in reduced parameter space. Notably, across all cases—cortical, striatal, hippocampal, thalamic, and cerebellar neurons—we found that neuromodulators such as acetylcholine, norepinephrine, serotonin, dopamine, and histamine shifted the excitability landscape of neurons in ways that could be compactly captured and visualized via principal component analysis (PCA) and phase plane analysis.

These findings are particularly interesting because neuromodulation dysfunction is often a key factor in brain disorders. For example, the two most common neurodegenerative diseases, Alzheimer’s and Parkinson’s, are associated with a decline in the cholinergic system and dopamine deficiency, respectively (***Chen et al., 2022; Latif et al., 2021***). In depression, imbalances in serotonin and norepinephrine contribute to symptoms (***Ressler and Nemeroff, 2000; Ruhé et al., 2007***), while schizophrenia is linked to dysfunction of the dopaminergic system (***Grace, 2016***).

In this context, a key insight from the striatal neuron modelling (Fig. 3) is relevant: dopamine modulation increases the separation between the parameter sets of direct-pathway (dSPNs) and indirect-pathway (iSPNs) spiny projection neurons. This suggests that dopamine not only modulates individual excitability but also amplifies the functional distinction between these two pathways. Hence, an intriguing future direction would be to test whether such divergence collapses in disease models—such as Parkinson’s—where dopaminergic input is impaired. If the dSPN and iSPN parameter distributions were to converge (resulting in a negative silhouette score in PCA), it might indicate a mechanistic biomarker for pathological states.

A conceptual strength of our work lies in our decision not to propose a single “best” model, but rather an ensemble of parameter sets that match the data within acceptable bounds (Supplementary Material 2). This is particularly important in cases where only a single electrophysiological trace is available, as is often the case with historical literature. Multiple parameter sets can reproduce the same trace but respond differently under varying inputs. Thus, having a diverse set of plausible models allows for more robust simulation of neurons at the network level and enables progressive refinement as more data becomes available. For instance, if additional traces with different input amplitudes were acquired, the ensemble could be re-ranked or pruned without restarting the fitting process. We used very broad sets of intervals for the starting AdEx parameters in our fitting. This is the worst possible case a scientist would encounter, and it still produces reasonably good results. Likely, in a more realistic scenario, a scientist has information suggesting narrower intervals for the starting parameters.

Additionally, we found it beneficial to plot all the phase planes for the top models during analysis. This allowed us to ensure that the selected representative models were not outliers but reflected the central tendency of the ensemble. This practice can reveal subtle qualitative differences in dynamics, such as the presence or absence of bistability or subthreshold oscillations, which may not be obvious from time-domain traces alone.

While the AdEx model is not designed to capture the detailed spike shape, we mitigated this limitation through visual inspection of the top-ranked models. This hybrid approach—quantitative fitting guided by a set of core features, followed by qualitative validation—ensured that selected models faithfully replicated the key dynamics of the source recordings. This methodological flexibility is particularly useful when raw data is unavailable and digitised traces from publications must suffice.

We also emphasise the technical contribution of this work: all modelling and simulations were conducted in Python, using custom scripts for AdEx simulation, feature extraction, parameter space investigation, dimensionality reduction, and visualisation. We deliberately chose not to use established simulation frameworks like NEST (***Gewaltig and Diesmann, 2007***) or Brian 2 (***Stimberg et al., 2019***), enabling full control and transparency over the parameter exploration and optimisation process, but also as a didactic choice. The decision to implement our own solvers and error functions allowed us to fine-tune the process, for example, by placinga greater penalty on poor spike-timing fits and incorporating additional features such as voltage at stimulus offset in adaptation-dominated traces.

One of the practical limitations we encountered during our parameter fitting was the lack of precise information regarding the holding current used to maintain neurons at a specific membrane potential in many experimental datasets. This uncertainty introduces variability in estimating the leak reversal potential (*E*_*L*_), an important parameter that reflects the resting membrane potential influenced by leak channels. The inability to confidently constrain *E*_*L*_ may introduce a degree of ambiguity in comparing fitted models, particularly when the holding current significantly differs across experimental setups.

Another point worth highlighting is the lack of correction for liquid junction potential in the source data we analysed. This systematic experimental oversight, common across many electro-physiological studies, can result in voltage offsets that propagate through the fitting and bias parameter estimations like *E*_*L*_ and *V*_*th*_. While not critical for relative comparisons within the same study, it does warrant caution when comparing across datasets or interpreting absolute parameter values.

Moreover, in many instances, the neuromodulator used in the experiments was a pharmacological agonist rather than the native molecule. For example, methacholine, a non-specific cholinergic agonist, was used instead of acetylcholine. While such substitutions are standard in experimental neuroscience, they must be kept in mind during interpretation since pharmacological agents may have broader or narrower receptor targets than endogenous ligands.

From a broader perspective, our work contributes to a growing body of literature that emphasises dynamical systems analysis as a bridge between experimental electrophysiology and computational modelling. While detailed Hodgkin-Huxley models offer biophysical fidelity, their complexity makes them less amenable to comparative analyses across neuron types and modulatory states. The AdEx model, by contrast, offers a tractable yet expressive framework that can be scaled to network simulations, subjected to mathematical analysis, and flexibly fitted to empirical data.

The implications of this work extend beyond single-neuron modelling. By providing compact descriptions of neuromodulatory effects, we pave the way for neuromodulation-aware network simulations, where different brain states (e.g., sleep vs. wakefulness, attention vs. drowsiness, motor execution vs. rest) can be modelled by adjusting parameter clusters in a biologically plausible way. Furthermore, our framework may support future studies on plasticity and learning, since neuromodulators are known to influence synaptic efficacy and circuit reconfiguration over time. The approach is extensible, generalizable, and particularly well-suited to the kinds of sparse, heterogeneous data that often characterise neuroscience research.

In conclusion, this study provides a structured and flexible methodology for capturing neuro-modulator effects on neural excitability across the central nervous system. We have provided a general approach to study how different families of neuromodulators affect neuronal dynamics. The dimensionality reduction analysis revealed that neuromodulation clusters parameters into distinct groups, allowing us to identify the most influential parameters for each neuromodulator. The joined phase plane analysis detailed how changes in those parameters affect the neuronal dynamics. By using interpretable parameters, clustering in feature space, and phase-plane analysis, we show how neuromodulation reshapes the landscape of neuronal dynamics.

## Methods and Materials

### Data collection and Curation

We systematically screened the available literature using targeted PubMed searches. For example, the search used for Acetylcholine was:

~~~
(acetylcholine[MeSH Terms] OR acetylcholine[Title/Abstract]) AND (electrophysiology[MeSH Terms] OR “patch clamp”[MeSH Terms] OR
“voltage clamp”[MeSH Terms] OR “current clamp”[MeSH Terms] OR
“intracellular recording”[MeSH Terms]) AND
(neuron[MeSH Terms] OR neuron*[Title/Abstract] OR “brain slice”[MeSH Terms])
~~~

This gives 1115 results, as of February 2025. We screened these results with the requirement that the original article must contain at least one voltage trace of control and <neuromodulator> conditions for the same recorded neuron.

Using this approach we were able to select voltage traces from five brain regions, representing seven neuronal types, recorded in control conditions and under the influence of five different neuromodulators. An overview of the recordings included in our dataset is provided in Table 1.

The selected publications did not provide raw electrophysiological data files, however the traces were available as figures. In order to retrieve the data, we extracted the voltage traces from images using the tool WebPlotDigitizer (***Rohatgi, 2010-2024***). While this approach is not ideal, it allowed us to reconstruct the traces with sufficient resolution for our analysis. All the extracted traces were saved in .csv format and are now available in the GitHub folder https://github.com/ Computational-NeuroPSI/Neuromodulation/tree/main/data/data_traces. A metadata file including also the holding voltage values and current protocols was also created and available in the same folder.

### Feature extraction

We developed Python scripts to load the `.csv` files and extract key electrophysiological features from each trace. These features, together with the associated current injection protocols, were saved in .json files which we then used to continue our modeling and analysis. As anticipated, we focused on features which allow us to describe the overall behaviour of the traces and are compatible with the level of detail we can retain. These key features are time to first, second, third and last spike (red bars 1, 2, 3, 4 in Fig. 1A and B), inverse of the first and last interspike interval (red bars 5 and 6, in Fig. 1A and B) and firing frequency.

From the metadata file, we read in the information about the current injection protocol. This includes the amplitude of the injected current (`curr`) expressed in pA as well as the temporal structure of the protocol: delay before stimulus onset (`stim_delay`), the time at which the stimulus ends (`stim_end`) and the total recording duration, all expressed in ms. From these we can easily compute the `stim_duration (stim_end - stim_delay)` and proceed with the extraction of the features.

The core mechanism underlying the extraction is the *detection of the action potentials*. We do that by identifying when the membrane potential crossed a fixed voltage threshold *V*_*th*_. This value depends on the neuron type and it is set by the user.

Specifically we initialize a `peak_index` list, iterate through the `trace_voltage` elements and if the following condition is verified, the corresponding index *i* is appended to `peak_index`:

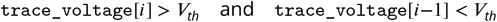

In other words, a spike is detected when the voltage trace crosses the voltage threshold “from below”.

From the `peak_index` list, we compute all the other features. The firing rate, freq, is computed as the number of spikes divided by the stimulus duration (in milliseconds) and expressed in Hertz:

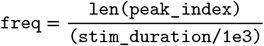

The time to first spike, `time_to_first_spike`, is the difference between the time at which the first peak is detected and the time at which the stimulation starts, we call the latter `stim_delay`. It is computed if at least one spike is detected:

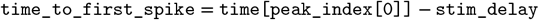

If at least two spikes are detected, we compute the `time_to_second_spike` and the inverse of the first interspike interval, `inv_first_ISI`, which corresponds to the inverse of the difference between the first and second spike time and it is expressed in Hertz:

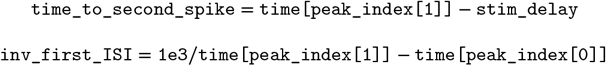

If exactly three spikes are detected, in an equivalent manner, we compute the `time_to_third_spike` and the `inv_last_ISI`:

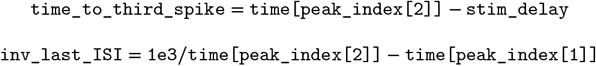

However, if more than three spikes are detected, the `inv_last_ISI` is computed as:

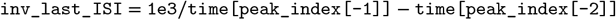

and the `time_to_last_spike` as:

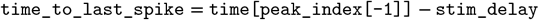

We selected the inverse of the interspike interval rather than the interval itself to express these values in units consistent with frequency, making them easier to visualise and interpret in the context of firing behaviour. Additionally, in cases where the traces exhibited strong spike-frequency adaptation (control DG neurons, Fig. 4 A; control thalamic relay neurons, Fig. 5A), we have also included the value of the membrane potential at the end of the stimulus as an additional feature. Specifically, if the last spike arrives in the first half of the stimulation, we compute it as the average of the voltage trace in the last 30 ms of the stimulation:

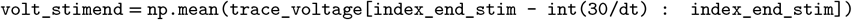

A jupyter notebook file exemplifying this process can be found here Neuromodulation/extracting_ features/Features_extraction.ipynb. We do not consider features which describe the spike shape because we do not have access to the raw data and also the AdEx model is not designed to reproduce detailed spike waveforms. However, we further refine the models that best match the extracted features by visually inspecting the spike shapes they generate and we prefer those that qualitatively resemble the recorded traces the best. This step is not mandatory, but it allows us to ensure that even if not explicitly fitted, the spike shapes are reasonably reproduced within the limits of the data itself and the model.

### Model simulation

A core mechanism of the algorithm is the simulation of the AdEx models, which consists in numerically solving the two coupled differential equations that describe the membrane potential and adaptation variable at each time step (same as Eq. 1):

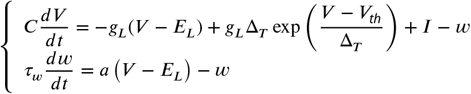

To do so, we implemented in Python the forward Euler numerical method. To handle the spike discontinuity, if the membrane potential *V* reaches or exceeds the peak voltage *V*_*peak*_, *V* is reset to *V*_*reset*_ and *w* is increased by *b*; then, for a refractory period (usually within 5 ms), both quantities are kept constant. While the forward Euler method is simple and less accurate than, for example, higher-order Runge-Kutta methods, its cheap computational cost and the smoothness of *V* and *w* before the peaks make it the preferred choice.

To provide a transparent and reproducible computational environment, we conducted all simulations using Python 3.9 and Jupyter Notebooks (***Kluyver et al., 2016***). The modeling environment also utilized specialized libraries for numerical simulations (Numpy, version 1.21.6, ***Harris et al., 2020***), data analysis (Scipy, version 1.8.1, ***Virtanen et al., 2020***), and visualization (Matplotlib, version 3.5.2, ***Hunter, 2007***).

To validate our implementation, we compared its solutions to those obtained using NEST and Brian 2, and confirmed that the results were qualitatively and quantitatively consistent. The models used for this comparison were initially taken from ***Naud et al., 2008*** (specifically, their Figure 4), which provided a useful starting point for testing the well-established simulators and our own implementation (Supplementary Fig. 1). As the project progressed, we also extended the comparison to include the models generated using our pipeline. Both versions of the comparison are available as separate notebooks here: https://github.com/Computational-NeuroPSI/Neuromodulation/tree/main/comparing_methods. It is important to note that each simulator uses its own conventions for units and proper unit conversion is therefore crucial to ensure consistency across implementations.

### Parameter exploration

The algorithm we developed takes as input the extracted features (reading them from the corresponding .json files) and the user-defined ranges for the AdEx model parameters. Additionally, the user can specify the number of models to generate. We explore the parameter space by sampling each parameter within its specified bounds. To efficiently sample the parameter space we use Sobol’s sequences via the python library `scipy.stats.qmc.Sobol` (***Joe and Kuo, 2008***). We preferred it to the standard pseudorandom sampling because Sobol sequences are quasi-random low discrepancy which ensures a more uniform and space-filling coverage of the parameter space. Since Sobol sequences are built recursively using binary representations, as a number of models we select a number which can be expressed as a power of 2. To maintain this structure, when selecting the number of best models we chose 16 (i.e. 2^4^). So, with the goal of reducing clustering and gaps in the high-dimensional space we want to investigate we selected this method, however the use can easily change how the space is sampled and still be able to continue with the analysis. Next step consists in running the newly obtained models and extracting the features. These features are the input of the error function which quantifies the difference between the data-derived and model-derived features.

### Model ranking

To evaluate how well each AdEx model reproduces the extracted features, we implemented an error function in Python (as part of the developed algorithm). This function, as anticipated, takes as input the values of each feature extracted from the data and the corresponding values extracted from the simulated models, and computes the relative error. It is defined as:

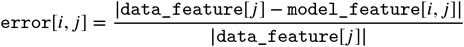

Where *i* indexes the models and *j* the features.

The total error for a given model is obtained by simply summing the relative errors across all features and all current protocols (if more than one is available). In cases where the model fails to “produce a feature” (for example, if it does not spike, many features can not be computed) we assign a penalty by adding +3 to the relative error for that specific feature. The value is arbitrary, it can of course be changed and the function can be personalized. Moreover, for example, in the cases where the traces to fit exhibited strong spike-frequency adaptation, the error in relation to the value of the membrane potential at the end of the stimulus was weighted more than those corresponding to other features.

Once the total error has been computed for each model, we rank all models in ascending order of error (by using `numpy.argsort`). We then store the top *n* models, parameter set and error values, in a . `json` to allow the user to further analyze them.

An example demonstrating both the parameter exploration and model ranking procedure can be found here: https://github.com/Computational-NeuroPSI/Neuromodulation/tree/main/exploring_parameters_space.

In many studies, it is common practice to use a single AdEx model per neuron type and condition. However, in this work, we take a different approach by selecting a set of models (Supplementary Fig. 2). During the parameter exploration and consequent fitting process, we often have access to only a single voltage trace, which may not be sufficient to constrain all the models in a way that ensures accurate reproduction of the f-I relation (when that curve is not available). By selecting the top *n* models from our fitting procedure, we aim to capture a broader range of plausible dynamics, and reduce the risk of using a single model that performs well on a single trace but might fail to generalize the neuron behaviour.

### Post-fitting analysis

We performed three main post-fitting analyses for each neuron type.

#### Comparing control and modulated AdEx parameters

We compared the AdEx parameters obtained for control and modulated conditions by examining the distribution of each parameter (Supplementary Material 2). Specifically, for each condition, we plotted the parameter values corresponding to the best 16 models and computed their mean and standard deviation. This allowed us to identify systematic shifts in the parameters space when neuromodulation was applied and also identify which parameters had a narrower range of variability. In this way we were able to spot the parameters which are more critical for capturing specific dynamics.

#### Principal Component Analysis

To visualize the structure of the parameter space and assess whether control and modulated models formed distinct groups, we performed a Principal Component Analysis on the best 16 models. The PCA revealed that models corresponding to different conditions formed clearly separate clusters in low-dimensional space and models fitted independently on traces corresponding to the same neuron and condition formed overlapped clusters. To quantify this separation, we computed the Silhouette score, which supported the reliability of the clustering. Both PCA and Silhouette score were computed using the scikit-learn library (***Pedregosa et al., 2011***).

#### Phase plane

Finally, since the AdEx model is described by two coupled differential equations, its dynamics can be easily visualized in a phase plane plot. In particular, we examine the nullclines (same as Eq. 3 and Eq. 4):

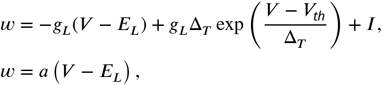

namely the curves where the derivatives of *V* and *w* are zero. The intersection of the two nullclines corresponds to fixed points. We investigated the stability of those by analyzing the eigenvalues of the Jacobian matrix obtained from linearizing the system around the fixed point. If all eigenvalues have negative real parts, the fixed point is stable (an attractor). If at least one eigenvalue has a positive real part, the fixed point is unstable. When the eigenvalues have both positive and negative real parts, the fixed point is a saddle point, exhibiting stable and unstable directions.

### Predicting the effect of a neuromodulator

The AdEx model allows us to predict the effect of neuromodulators across species. For example, we used rodent data to predict the effect of dopamine on human cortical neurons (Fig. 8). To do so, two strategies were implemented. For the first strategy, *absolute value strategy*, we computed a *modulation ratio* for each parameter *i* as:

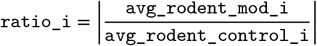

where `avg_rodent_mod_i` and `avg_rodent_control_i` represent the mean value of the *i*-th parameter under modulation and in control conditions, respectively. Then we apply these relative changes observed in rodent to the average human control values:

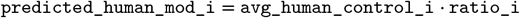

for each parameter *i* of the AdEx model.

However, if the corresponding control values of the two species do not have the same sign, the signed change is not preserved. This is for example the case of the parameter *a* (Fig. 8C and Supplementary Fig. 4A).

In order to take that into account, a second strategy was implemented. In this case, the relative change for the *j*-th parameter which has different signs for the two species (in this case it is the parameter *a*) is computed as:

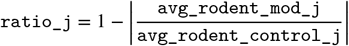

Then, analogously to the previous case, the *j*-th predicted value is computed as:

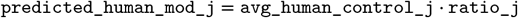

This approach reflects the same direction of change even if the baseline values have opposite signs (Supplementary Fig. 4C) and for this reason we refer to this as *signed value strategy*. Both strategies originate comparable results in terms of predicted dynamics, while the increase in firing frequency is more pronounced in the second case (Supplementary Fig. 4B and D).

A notebook exemplifying these approaches can be found in the GitHub folder.

## Software and code availability

To ensure transparency and facilitate replication, we made our code (fully written in Python) and software publicly available on GitHub: https://github.com/Computational-NeuroPSI/Neuromodulation. The repository contains a complete pipeline for analyzing neuromodulatory effects on neuronal excitability using the AdEx model. It includes:

- Curated electrophysiological data derived from figures in published papers,
- Feature extraction tools to process the voltage traces
- AdEx simulation code using our implementation based on the forward Euler numerical method, NEST and Brian 2,
- Parameter space exploration and filtering of best models,
- Post-fitting analyses including parameter distribution comparisons, PCA and phase-plane plots,
- Prediction of neuromodulatory effects in species lacking direct data,
- Example scripts and notebooks to reproduce results and visualize model behavior.

Simulations were performed on a Linux-based desktop computer equipped with an AMD^®^ Ryzen™ 9 3900X 12-core (24-thread) processor and 64 GB of RAM.

## Acknowledgments

Research supported by CNRS, Agence Nationale de la Recherche (BrainAct and FLAG-ERA JTC ImpactCom projects), and the European Union (Human Brain Project H2020-945539 and Virtual Brain Twin project 101137289).

## Author contributions

D.G. contributed to conceptualization, data curation, development of analytic tools, formal analysis, and wrote the manuscript.

I.C. contributed to conceptualization, data curation, development of analytic tools, formal analysis; performed the simulations, analysed the results; and wrote the manuscript.

A.D. contributed to conceptualization, supervised the project.

