## Supplementary Material for "Neuromodulators control neuronal dynamics through feature space reshaping"

#### Supplementary Material 1

In this section we report the Supplementary Figures:

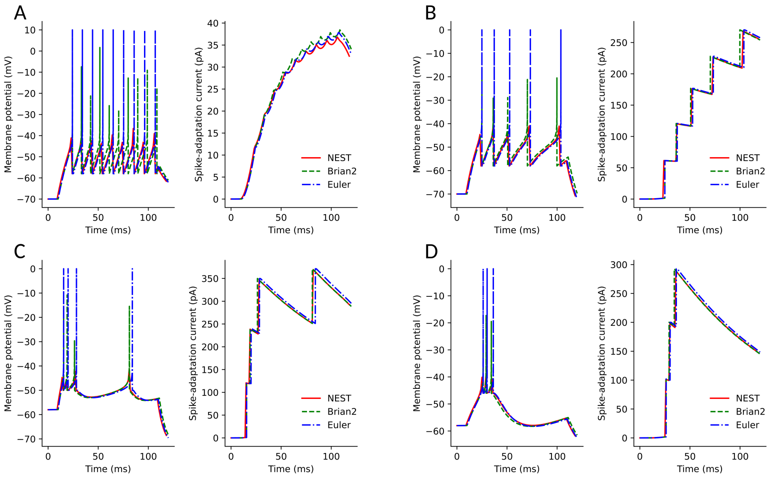


**Supplementary Figure 1. Comparing NEST, Brian 2 and our implementation when using models from *Naud et al., 2008*.** Comparison of three different simulation environments for solving Adaptive Exponential Integrate-and-fire (AdEx) neuron models: NEST (red, solid), Brian 2 (green, dashed), and our own implementation based on the Euler method (blue, dash-dot) when using the models from Figure 4 A-D in *Naud et al., 2008* which correspond to panel A-D, respectively. The lines represent the membrane potential (left of each panel) and spike-adaptation current (right of each panel) obtained by numerically solving the two equations describing the AdEx. Both NEST and Brian 2 do not draw the full spike, and in case of Brian 2, the slight temporal mismatch arises from the use of its default refractory period.

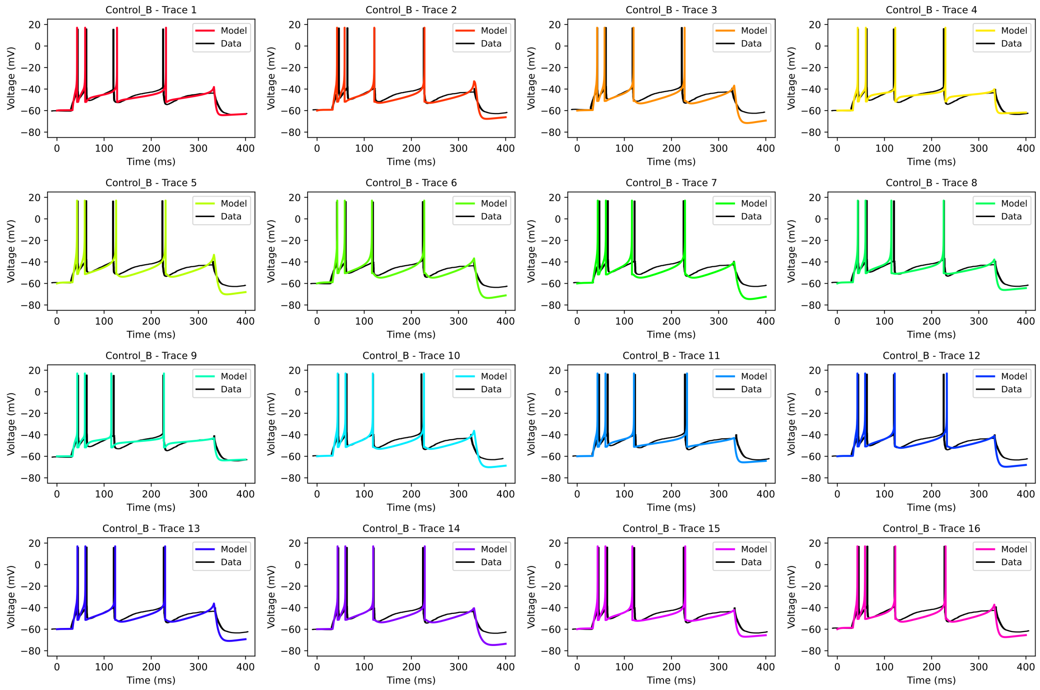


**Supplementary Figure 2. Best $16$ cortical pyramidal neuron models in control condition.** Comparison between the $16$ best models (colored lines) and experimental data (back line) extracted from Fig. 1 B in ***McCormick and Williamson, 1989***. Each subplot shows the membrane potential response for a single model and the experimental voltage recording.

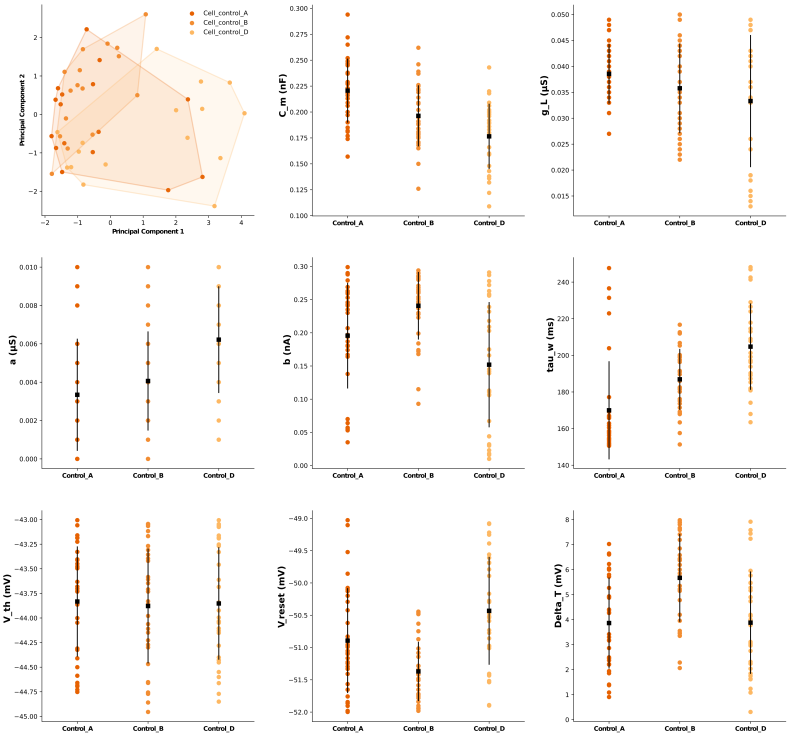


**Supplementary Figure 3. Comparison of AdEx model parameters fitted to three electrophysiological traces from the same layer III human cortical neuron.** (Top left) Principal Component Analysis (PCA) of AdEx models independently fitted to three current-clamp traces from ***McCormick and Williamson, 1989*** (their Fig. 1 A, B and D) recorded from the same layer III human cortical neuron using the same current injection protocol. These traces differ by eye, however the parameters of the models they originate are similar. To quantify this similarity, we run the PCA which shows strong overlap between the three clusters (each representing the models obtained from one of the traces), indicating similarity in their underlying parameter structure. Indeed the corresponding silhouette score has a negative value ($\sim -0.028$), indicating indeed poor cluster separation. (Remaining panels) Distribution of selected AdEx parameters (each subplot represents one parameter distribution) where the columns represent the models fitted to one of the three traces, the dots represent single models, while black squares and bars indicate the mean and standard deviation, respectively.

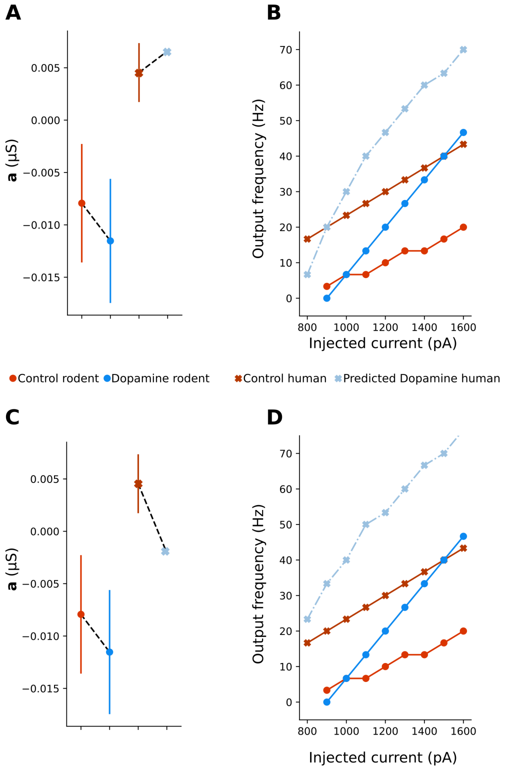


**Supplementary Figure 4. The effect of dopaminergic modulation on the subthreshold adaptation parameter.** The parameter $a$ (of the AdEx model) exhibits opposite baseline values in rodents and humans. In rodent control models $a$ is negative (red circle in A and C represents the average) and it further decreases under dopaminergic modulation (blue circle in A and C represents the average). In contrast, human control models (whose average is represented with a red x mark in A and C) show a positive $a$. This leads to two possible strategies for predicting the dopaminergic effect on humans: (A) by increasing the absolute value of $a$ and (C) by applying signed decrease. Panels A and C show the resulting changes in $a$ for both strategies, while panels (B) and (D) compare the corresponding output frequency–current curves. Both approaches produce comparable predicted firing dynamics, although the signed decrease (C–D) results in a more pronounced increase in firing frequency.

#### Supplementary Material 2

In this section, we provide a comparative analysis of AdEx parameters across control and neuromodulated conditions for seven neuron types: human cortical pyramidal neurons, striatal direct and indirect projection neurons, dentate gyrus neurons, thalamocortical projecting neurons, thalamic reticular neurons, and cerebellar Purkinje neurons. For each neuron type and condition, we display the distributions of the AdEx parameters corresponding to the top $16$ models, alongside their means and standard deviations.

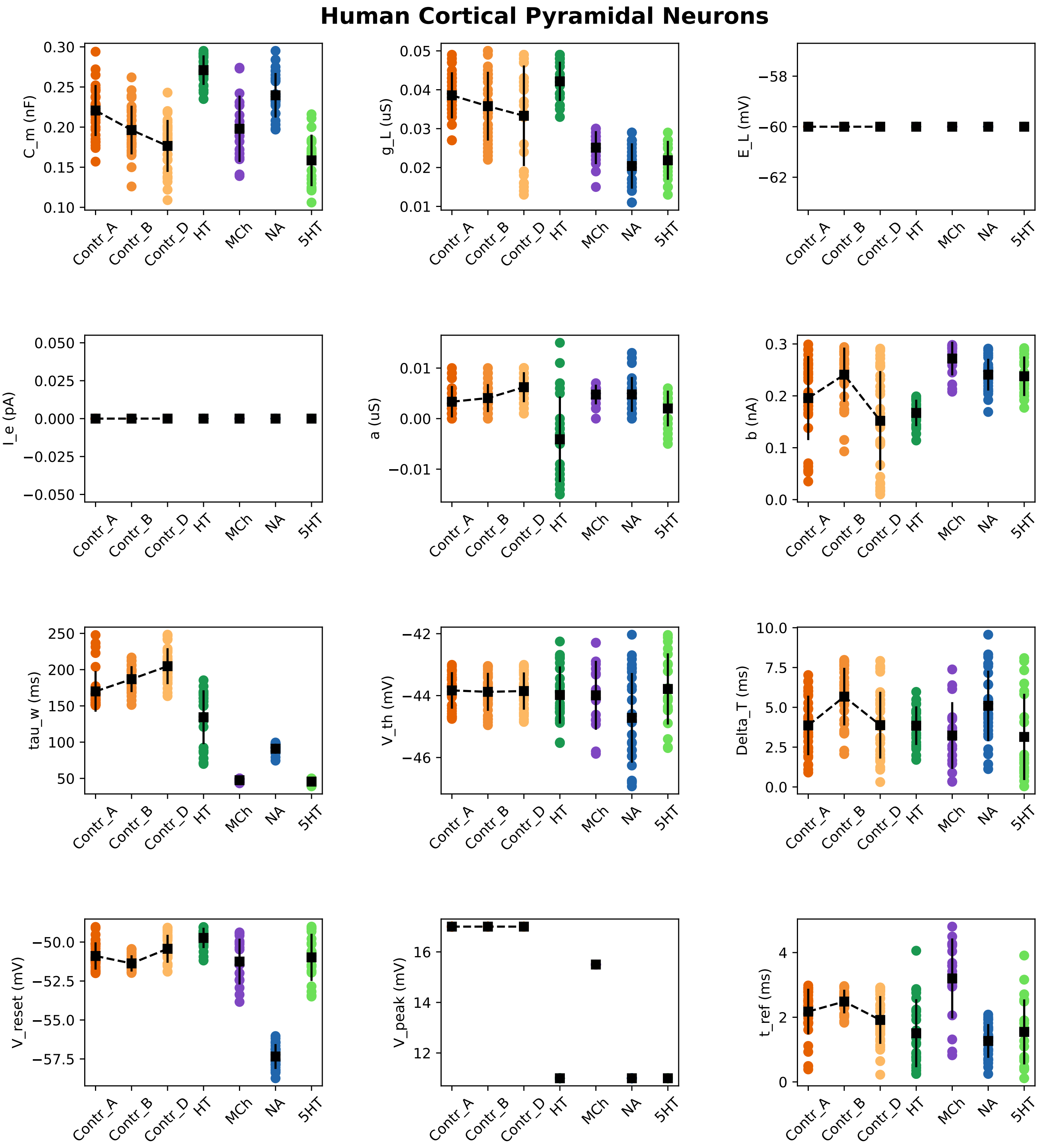
Human Cortical Pyramidal Neurons
C_m (nF)
g_L (uS)
E_L (mV)
I_e (pA)
a (uS)
b (nA)
tau_w (ms)
V_th (mV)
Delta_T (ms)
V_reset (mV)
V_peak (mV)
t_ref (ms)
Contr_A
Contr_B
Contr_D
HT
MCh
NA
5HT

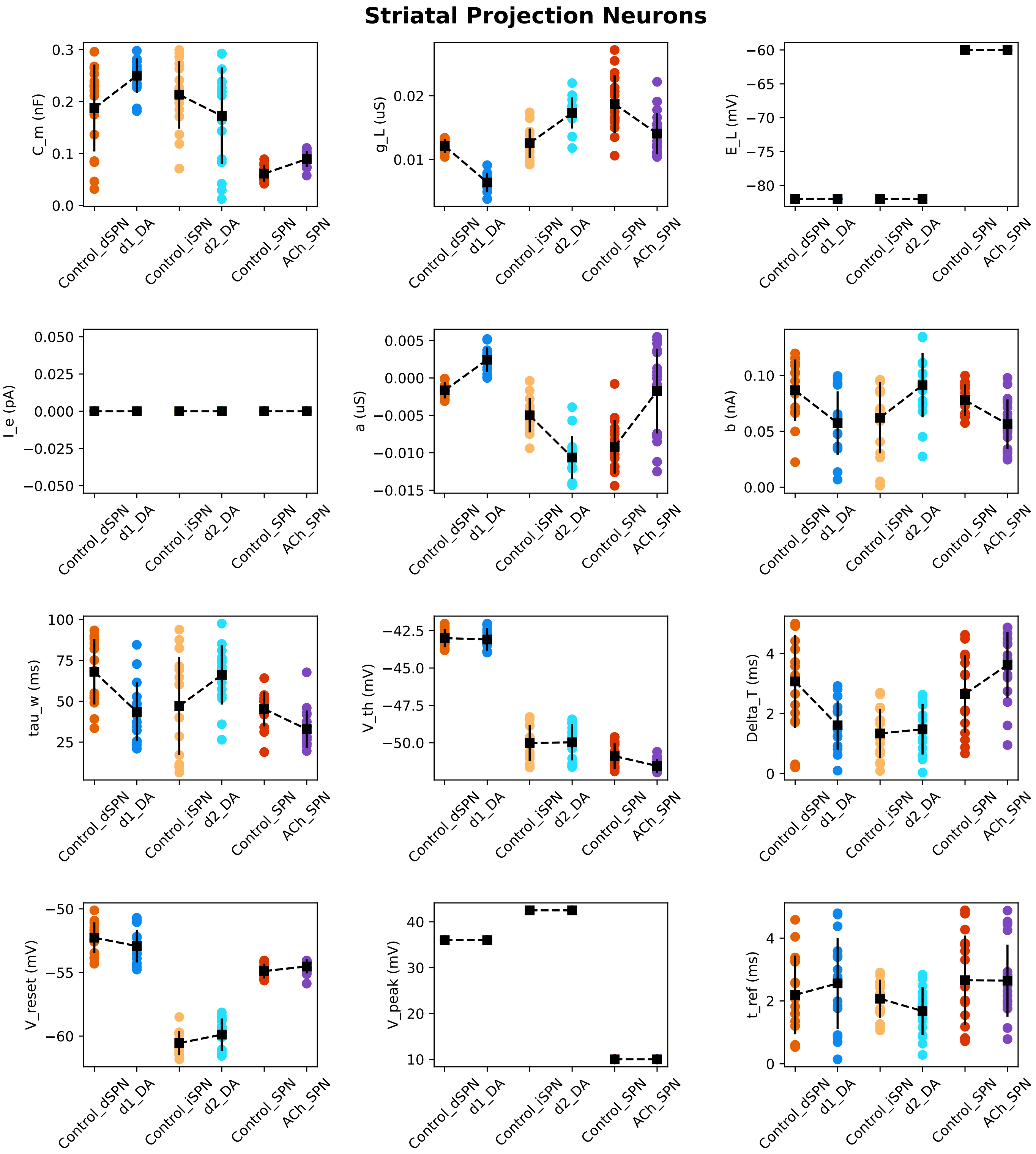
Striatal Projection Neurons
C_m (nF)
g_L (uS)
E_L (mV)
I_e (pA)
a (uS)
b (nA)
tau_w (ms)
V_th (mV)
Delta_T (ms)
V_reset (mV)
V_peak (mV)
t_ref (ms)
Control_dSPN
d1_DA
Control_iSPN
d2_DA
Control_SPN
ACh_SPN

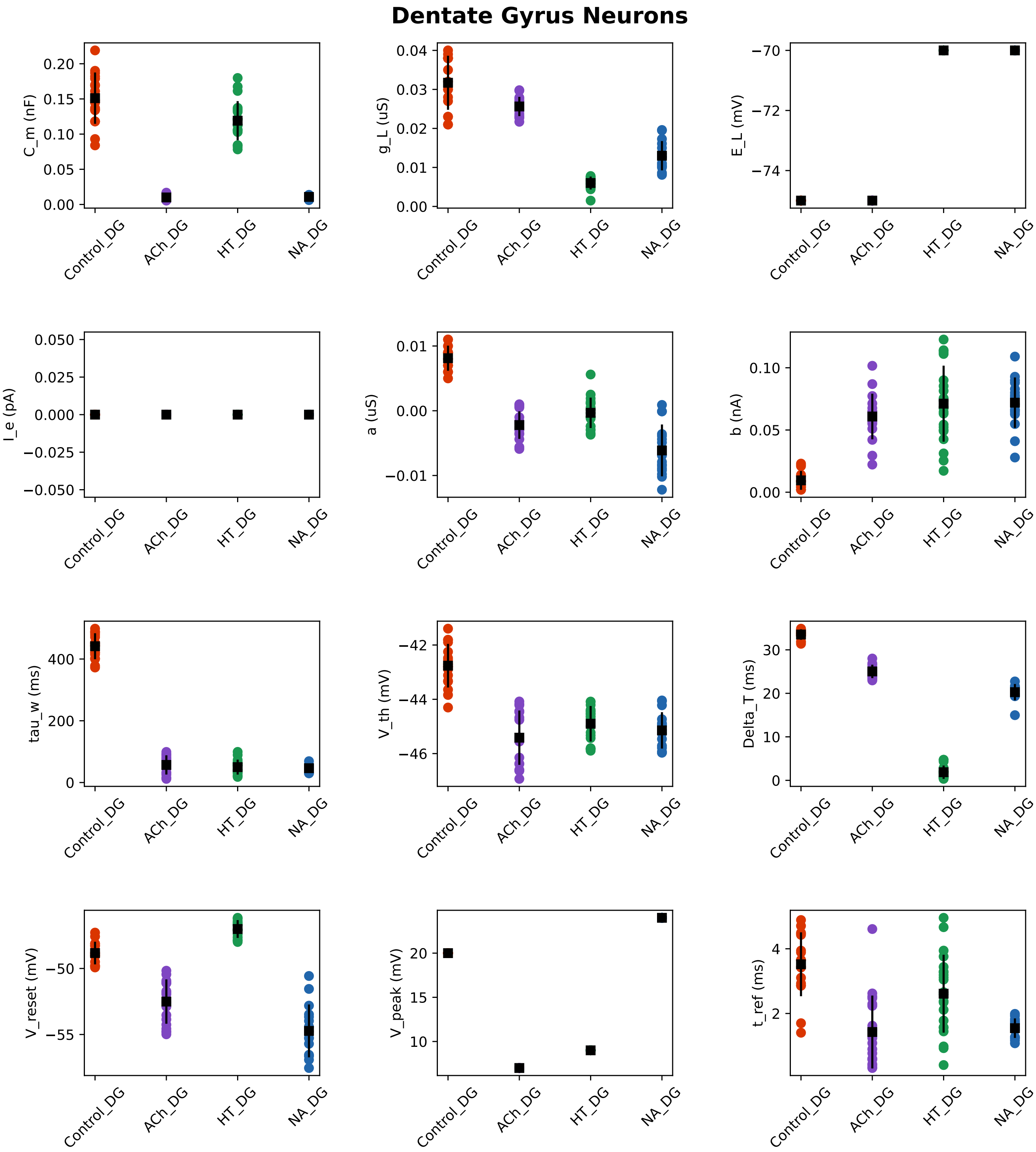
Dentate Gyrus Neurons
C_m (nF)
g_L (uS)
E_L (mV)
I_e (pA)
a (uS)
b (nA)
tau_w (ms)
V_th (mV)
Delta_T (ms)
V_reset (mV)
V_peak (mV)
t_ref (ms)
Control_DG
ACh_DG
HT_DG
NA_DG

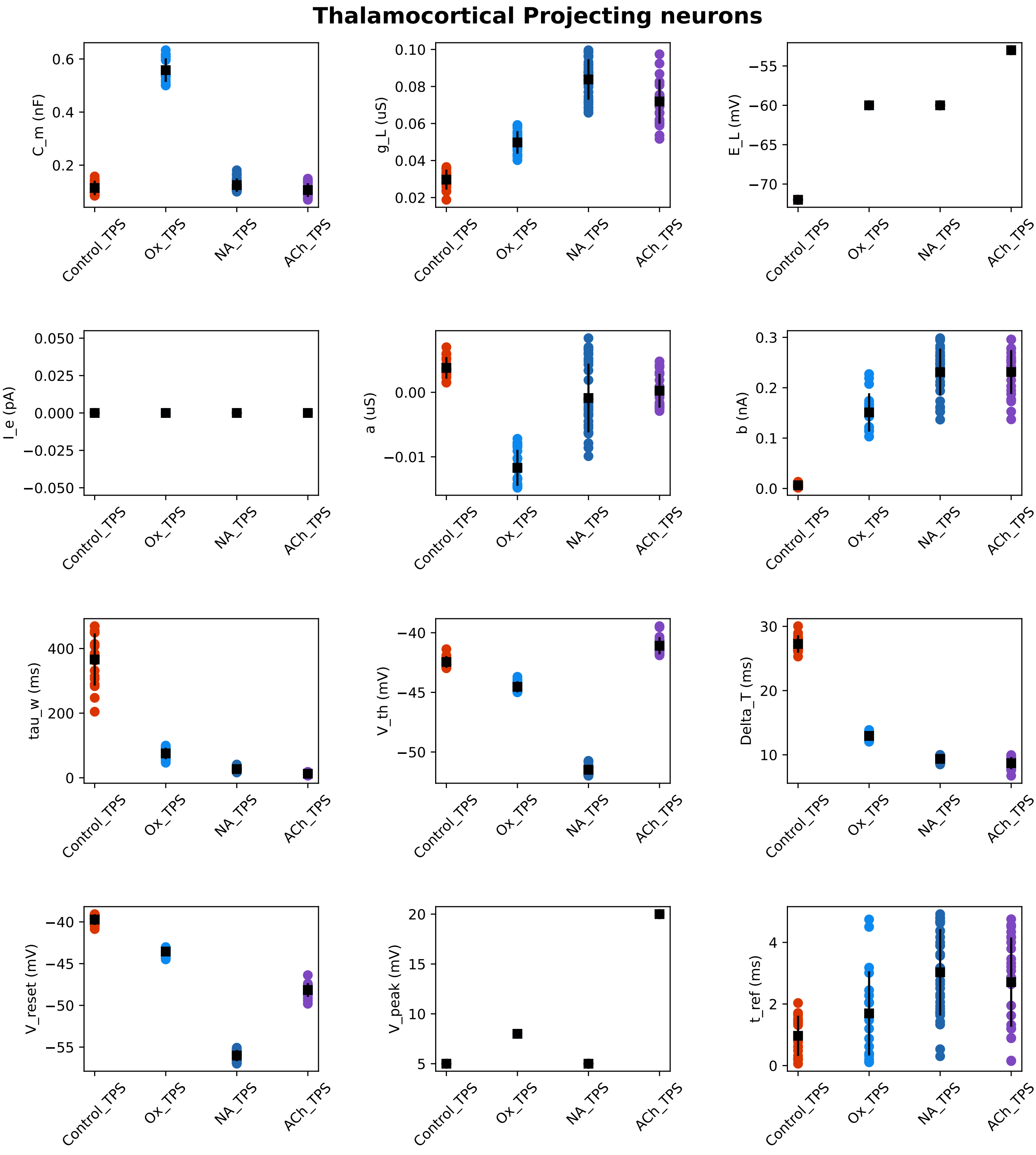
Thalamocortical Projecting neurons
C_m (nF)
g_L (uS)
E_L (mV)
I_e (pA)
a (uS)
b (nA)
tau_w (ms)
V_th (mV)
Delta_T (ms)
V_reset (mV)
V_peak (mV)
t_ref (ms)
Control_TPS
Ox_TPS
NA_TPS
ACh_TPS

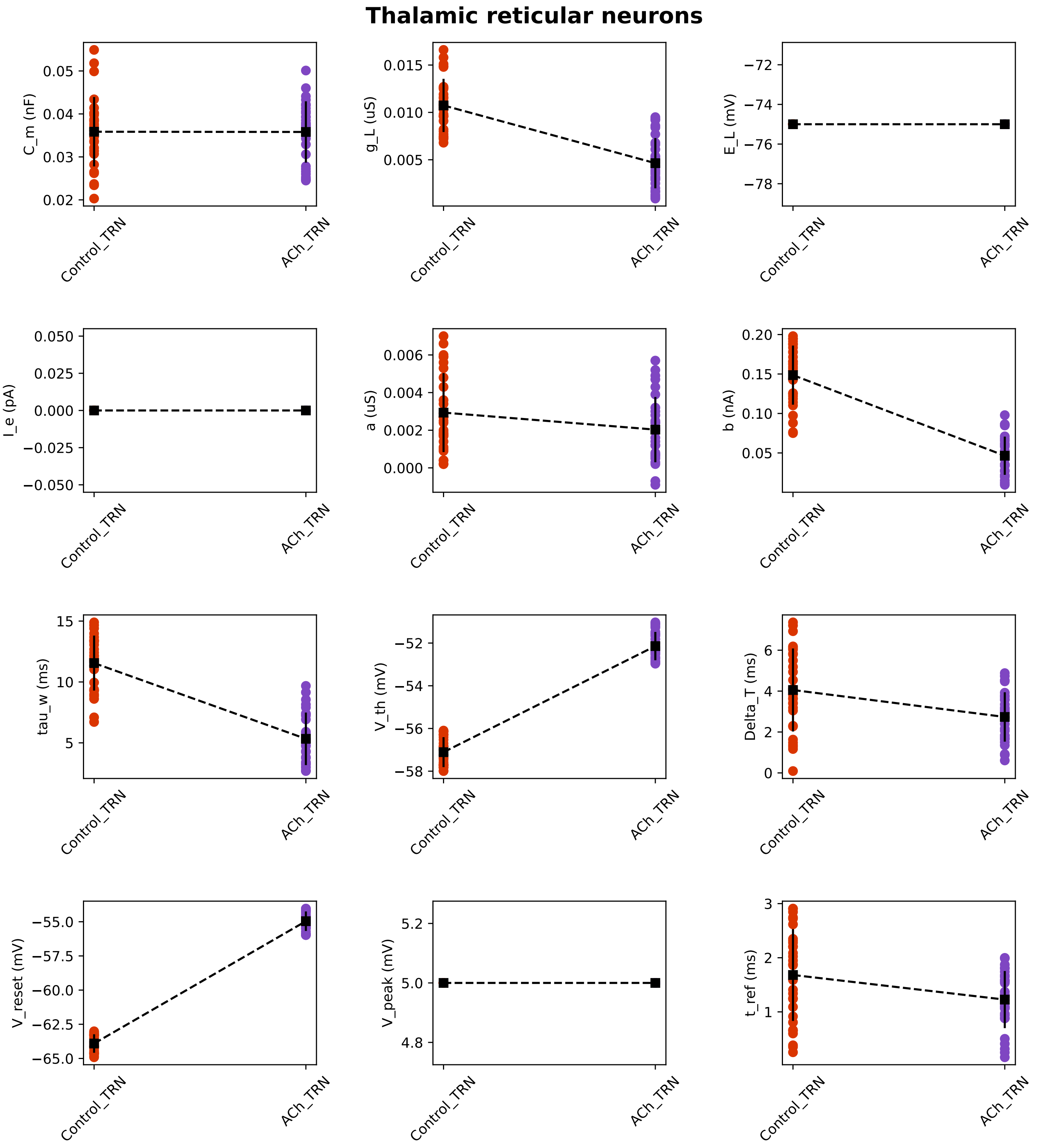
Thalamic reticular neurons
C_m (nF)
g_L (uS)
E_L (mV)
I_e (pA)
a (uS)
b (nA)
tau_w (ms)
V_th (mV)
Delta_T (ms)
V_reset (mV)
V_peak (mV)
t_ref (ms)
Control_TRN
ACh_TRN

### Cerebellar Purkinje neurons

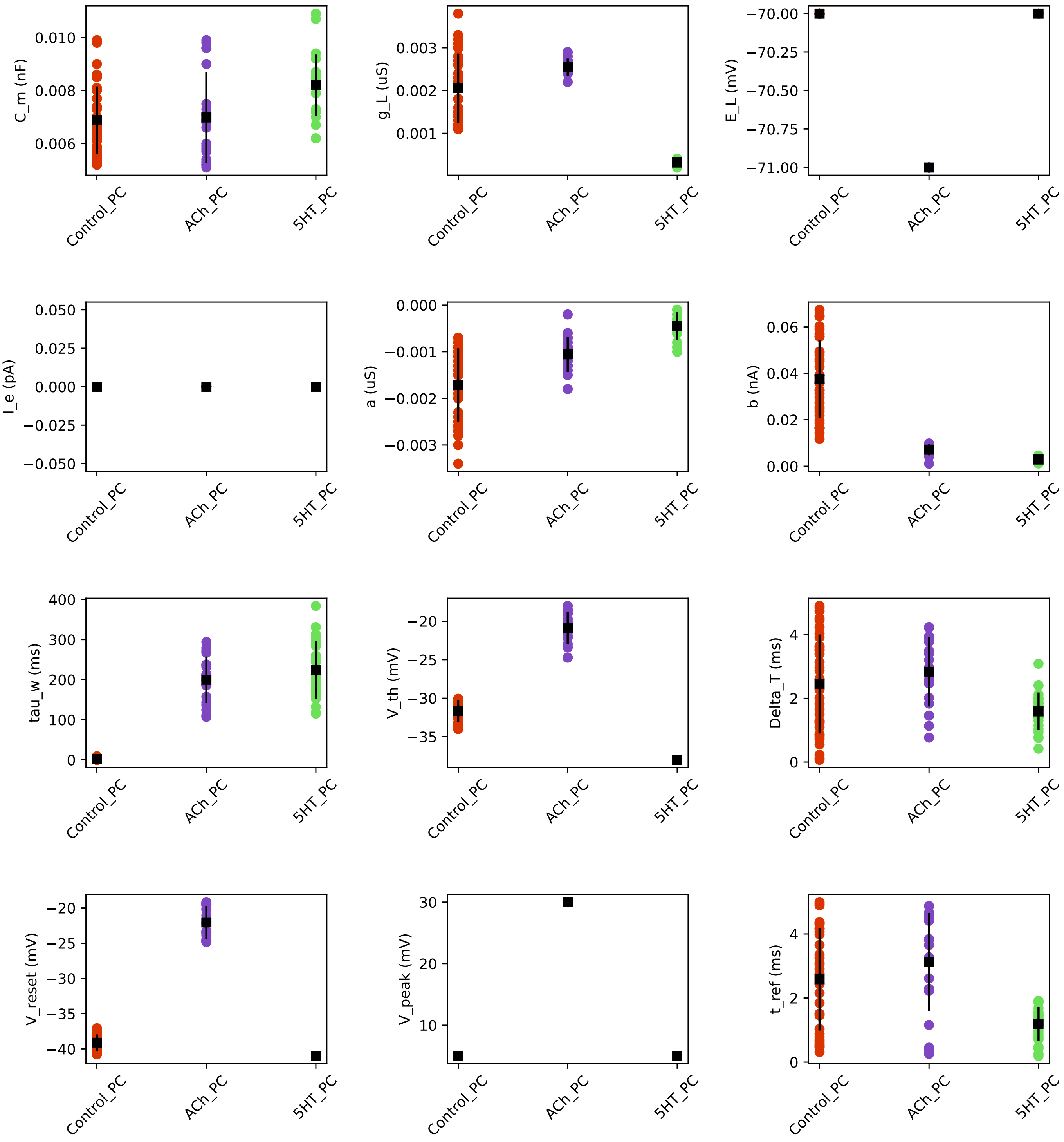
